# Cortical assembloids support the development of fast-spiking human PVALB+ cortical interneurons and uncover schizophrenia-associated defects

**DOI:** 10.1101/2024.11.26.624368

**Authors:** Ryan M. Walsh, Gregg W. Crabtree, Kriti Kalpana, Luz Jubierre, So Yeon Koo, Gabriele Ciceri, Joseph A. Gogos, Ilya Kruglikov, Lorenz Studer

## Abstract

Disruption of parvalbumin positive (PVALB+) cortical interneurons is implicated in the pathogenesis of schizophrenia. However, how these defects emerge during brain development remains poorly understood. The protracted maturation of these cells during postnatal life has made their derivation from human pluripotent stem cells (hPSCs) extremely difficult, precluding hPSC-based disease modeling of their role in neuropsychiatric disease. Here we present a cortical assembloid system that supports the development of PVALB+ cortical interneurons which match the molecular profiles of primary PVALB+ interneurons and display their distinctive electrophysiological features. Further, we characterized cortical interneuron development in a series of CRISPR-generated isogenic structural variants associated with schizophrenia and identified variant-specific phenotypes affecting cortical interneuron migration and the molecular profile of PVALB+ cortical interneurons. These findings offer plausible mechanisms on how the disruption of cortical interneuron development may impact schizophrenia risk and provide the first human experimental platform to study of PVALB+ cortical interneurons.

## Introduction

Schizophrenia is a debilitating disorder associated with a diverse group of positive, negative, and cognitive symptoms. Though schizophrenia is typically diagnosed in adolescence, multiple lines of evidence suggest an early neurodevelopmental origin of the disease^1,2^. Of note, the cognitive deficits associated with schizophrenia are commonly observable in childhood, preceding the onset of positive symptoms and schizophrenia diagnosis. EEG and MEG studies in human patients performing cognitive tasks have identified defects consistent with a disruption of the function of fast-spiking parvalbumin (PVALB+) cortical interneurons^3–8^. Indeed, disruptions in GABAergic cortical interneurons are frequently observed in postmortem analysis of schizophrenia patients^9–13^. However, these observations in human patients were made after the onset of symptoms; as such, any developmental drivers including disease-related disruptions in cortical interneurons remain unknown. Therefore, a human system that replicates early cortical interneuron development in the context of schizophrenia could provide a valuable look into those critical early stages of the disease.

Estimates of the heritability of schizophrenia range from 65-81%^14^, suggesting it is a highly heritable disorder. The genetics of the disease, however, is complex, and in many patients, the pathology is likely the result of the combined effect of multiple low-risk common variants^15–20^. Interestingly, there is strong evidence implicating large effect rare variants in the genetics of the disorder^21–25^. Among the most consistently schizophrenia-associated rare variants are several large genomic structural variants (SVs). While these SVs represent a comparatively small percentage of schizophrenia patients, it is likely that the study of such cases in a human system could provide broad mechanistic insights that may apply to the disorder as a whole.

During development, cortical interneurons are produced primarily within the ganglionic eminences of the ventral forebrain and undergo a tangential migration into the dorsal forebrain where they integrate into circuits with the locally born excitatory neurons of the cortex^26^. The majority of cortical interneurons (60- 70%) are produced within the medial ganglionic eminence (MGE), a region that gives rise to both PVALB+ and somatostatin (SST+) subtypes with a smaller populations of (∼30%) VIP+ and SCGN+ interneurons derived from the caudal ganglionic eminence (CGE)^26,27^. Recent evidence suggests an additional locus of interneuron generation in the human dorsal cortex, yet this population appears to be molecularly similar to CGE derived interneurons^28^. Disruptions in cortical interneuron tangential migration and integration into the cortex have been linked to neurologic disorders in animal studies and in some human studies ^29–34^, however it is not clear what role disruptions in this process may play in schizophrenia. Indeed, animal models investigating schizophrenia associated genes and SVs have found evidence of disrupted migration in neurons and specifically in cortical interneurons^30,31,35^. However, patient data has been less clear, with some studies suggesting a reduction in cortical interneuron density^36–41^ while a number of others observe decreases in cortical GABAergic gene expression without a reduction in cortical interneuron density^10–13^. The process of cortical interneuron tangential migration has been recently modeled in cortical assembloid culture^42–44^, providing an attractive system for investigating whether disruption of cortical interneuron tangential migration does indeed represent a risk factor in schizophrenia.

Previous studies have employed patient-specific induced pluripotent stem cell (iPSC)-derived neuronal and neural organoid models to study aspects of schizophrenia *in vitro*, with different groups reporting decreased progenitor survival and neurogenesis defects^45^, mitochondrial defects^46–50^, reduced expression of neuron developmental factors^51^, altered Na^+^ channel function^52^, and reduced connectivity or synaptic defects^49,53,54^ (further reviewed in^55^). While these studies have provided interesting insights into possible molecular defects involved in schizophrenia, it is difficult to address the impact of confounding factors like genetic background and, in some cases, the precise identity and type of the neurons being assessed. Given that defects in schizophrenia have been reported in specific neuronal types and subtypes, most notably in the PVALB+ interneuron population in the human cortex, the ability to derive and mark specific classes of neurons *in vitro* may be critical for generating reliable models of schizophrenia. We and others have demonstrated expression of cortical interneuron markers as well as PVALB expression in our *in vitro* directed differentiation systems^42–44,56–60^. While the ability to study fast-spiking PVALB+ cortical interneurons *in vitro* is an exciting prospect, PVALB is also expressed in populations of GABAergic neurons other than cortical interneurons, such as the ventral pallidum, hypothalamus, thalamic reticular nucleus and midbrain^61–65^. Furthermore, in vivo, PVALB expression in fast spiking cortical occurs largely during postnatal and adolescent stages, a time frame difficult to capture in current hPSC culture system. A more comprehensive assessment of PVALB interneuron identity would require, in addition to the developmental MGE origin with co-expression of FOXG1 and NKX2.1, the presence of fast-spiking electrophysiology features, and the persistent expression of the cortical interneuron marker LHX6 within PVALB positive interneurons.

Here we established a platform that enables the study of cortical interneuron development, including the derivation of bona fide PVALB+ cortical interneurons, and their role in schizophrenia pathogenesis. We created a cortical assembloid protocol that, similar to previous forebrain assembloids, generates GABAergic cortical interneurons and glutamatergic neurons in separately patterned ventral and dorsal forebrain organoids prior to their aggregation. However, our culture conditions support the emergence of LHX6+/PVALB+ cortical interneurons within 120 days of culture, without the need of transgene expression or *in vivo* cell transplantation. The identity of the PVALB+ human cortical interneurons was characterized by scRNA-seq and by using real time genetic reporters. Electrophysiological recordings in PVALB+ neurons showed firing patterns consistent with *bona fide* PVALB+ cortical interneurons, indicating a level of maturity not previously reported *in vitro*. Cortical assembloids also showed robust activity organized into complex neural network oscillations with power spectrum peaks in all major oscillatory bands including γ. We further generated a set of isogenic schizophrenia-associated SVs and characterized their effect on tangential migration within the same assembloids as their isogenic controls. We found that a hemizygous deletion of the 22q11.2 locus, which is both the most common SV in humans and the SV that confers the greatest risk for developing schizophrenia, led to defects in cortical interneuron migration and a reduction of 22q11.2-deleted cortical interneurons within the dorsal forebrain portion of cortical assembloids. We further observed a disruption in mitochondrial gene expression and function that serves as a plausible explanation for the tangential migration defect in 22q11.2 deletion interneurons. Finally, scRNA-seq analysis of PVALB+ cortical interneurons carrying hemizygous 15q13.3 deletion, showed likely disease-related changes in cadherin and protocadherin expression compared to their isogenic counterparts. These changes were highly specific and not seen in the related SST+ cortical interneuron population. Our study presents a platform suitable to determine SV- specific defects related to schizophrenia risk at distinct stages of cortical interneuron development.

## Results

### Generation of isogenic structural variant lines associated with psychiatric disorders

To study aspects of cortical interneuron and forebrain development that may contribute to psychiatric disease, we selected a set of 4 disease-associated structural variants (SVs). Additionally, each of these 4 SVs has a reported cortical interneurons defect in animal studies (Suppl. Table 1), making them prime candidates to study disease-relevant cortical interneuron dysfunction in a human system. Two of these, the 22q11.2 deletion and 15q13.3 deletion, have demonstrated an association with schizophrenia and other psychiatric disorders with an odds ratio of 11.5 or greater^21,66–68^ (Suppl. Table 1). The third of these SVs, is a 400kb deletion in chromosome 2q34 and was identified as a *de novo* mutation in a single patient with childhood onset schizophrenia^22^. However, given evidence linking such *de novo* SVs to schizophrenia and the fact that this deletion encompasses a portion of *ERBB4*, a gene required for multiple stages of PVALB cortical interneuron development^35,69,70^, we decided to include it in our study. The 22q11.2 deletion and 15q13.3 deletion arise through non-allelic homologous recombination (NAHR) between flanking segmental duplications, and this process can be induced *in vitro* by using CRISPR/Cas9 with a sole sgRNA targeting a homologous sequence in these segmental duplications^71^. We found that a single sgRNA targeting the segmental duplication was sufficient to induce both large scale genomic deletions and duplications (Fig. 1A). As might be expected, the efficiency of inducing these NAHR-mediated genetic lesions was dependent upon the size of the SV, with 6.7% of clones analyzed carrying the 1.6Mb 15q13.3 deletion compared to only 3.4% harboring the 2.5Mb 22q11.2 deletion (Fig. 1B). The *de novo* 2q34 deletion, on the other hand, lacks any flanking segmental duplications and therefore is likely the result of non-homologous end joining (NHEJ). As such it was generated *in vitro* with CRISPR/Cas9 and a pair of sgRNAs targeting either side of the observed SV (Fig. 1A&B).

**Figure 1.**
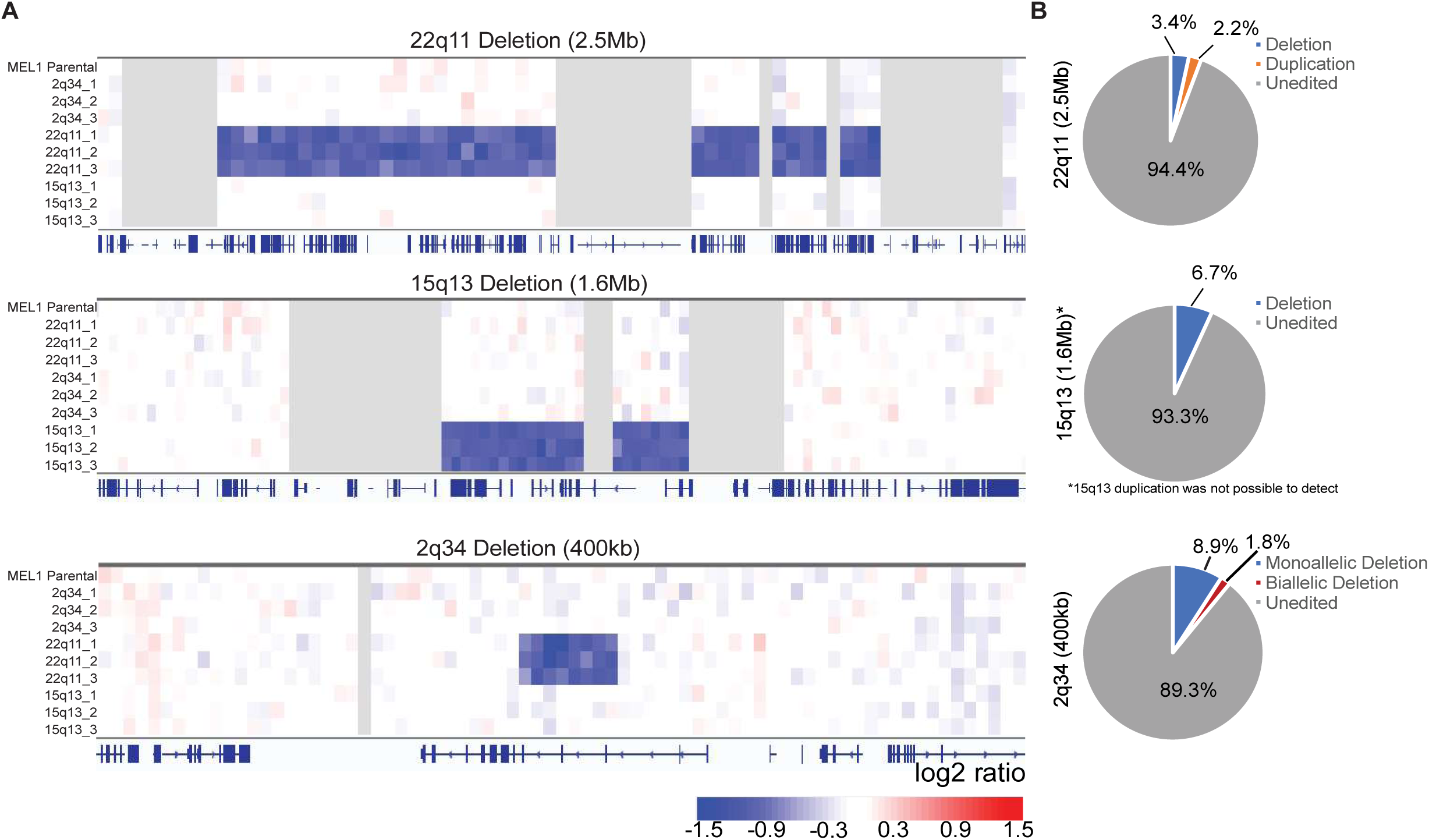
Generation of isogenic lines carrying schizophrenia associated structural variants. **A)** Data from shallow whole genome sequencing from the parental MEL-1 line and the 3 SV clones each to be used in this study. The genomic region containing each SV is displayed. **B)** Efficiency of SV generation by NAHR and NHEJ as determined by screening with TaqMan copy number assay, or genomic PCR for the 15q13 del. Due to the assay setup for 15q13, it was not possible to detect duplications and therefore only deletions are displayed.

### Patterning and aggregation of dorsal and ventral forebrain organoids

The neocortex comprises glutamatergic neurons and GABAergic cortical interneurons, which are two broad classes of neurons with distinct developmental origins. While the glutamatergic neurons are generated locally within the developing cortex, cortical interneurons are primarily born in the ventral region of the forebrain and must tangentially migrate to reach and integrate in the pallium^26^. Recent studies have demonstrated the feasibility of using directed differentiation to pattern separate dorsal and ventral forebrain-like organoids and subsequently fuse them into cortical “assembloids” containing both cortical glutamatergic neurons and GABAergic cortical interneurons^42,43,60,72^. We have previously achieved efficient forebrain induction using a combination of dual-SMAD inhibition and early WNT inhibition in both monolayer^56,73^ and 3D cultures^74^. This differentiation paradigm (Fig. 2A, top) reliably produces dorsal forebrain (dFB) organoids with a pattern of gene expression and cellular organization reminiscent of the developing telencephalon, without requiring Matrigel coating. Notably, the resulting cells broadly express the forebrain marker FOXG1 and contain ventricular-like zones of progenitors expressing the dFB progenitor marker PAX6 surrounded by a subventricular-like zone containing TBR2 expressing intermediate progenitors (Fig. 2B). Additionally, dFB organoids display a cortical-like laminar distribution of neurons with deep layer neuron markers expressed adjacent to progenitor zones and superficial markers expressed distally (Fig. 2B).

**Figure 2.**
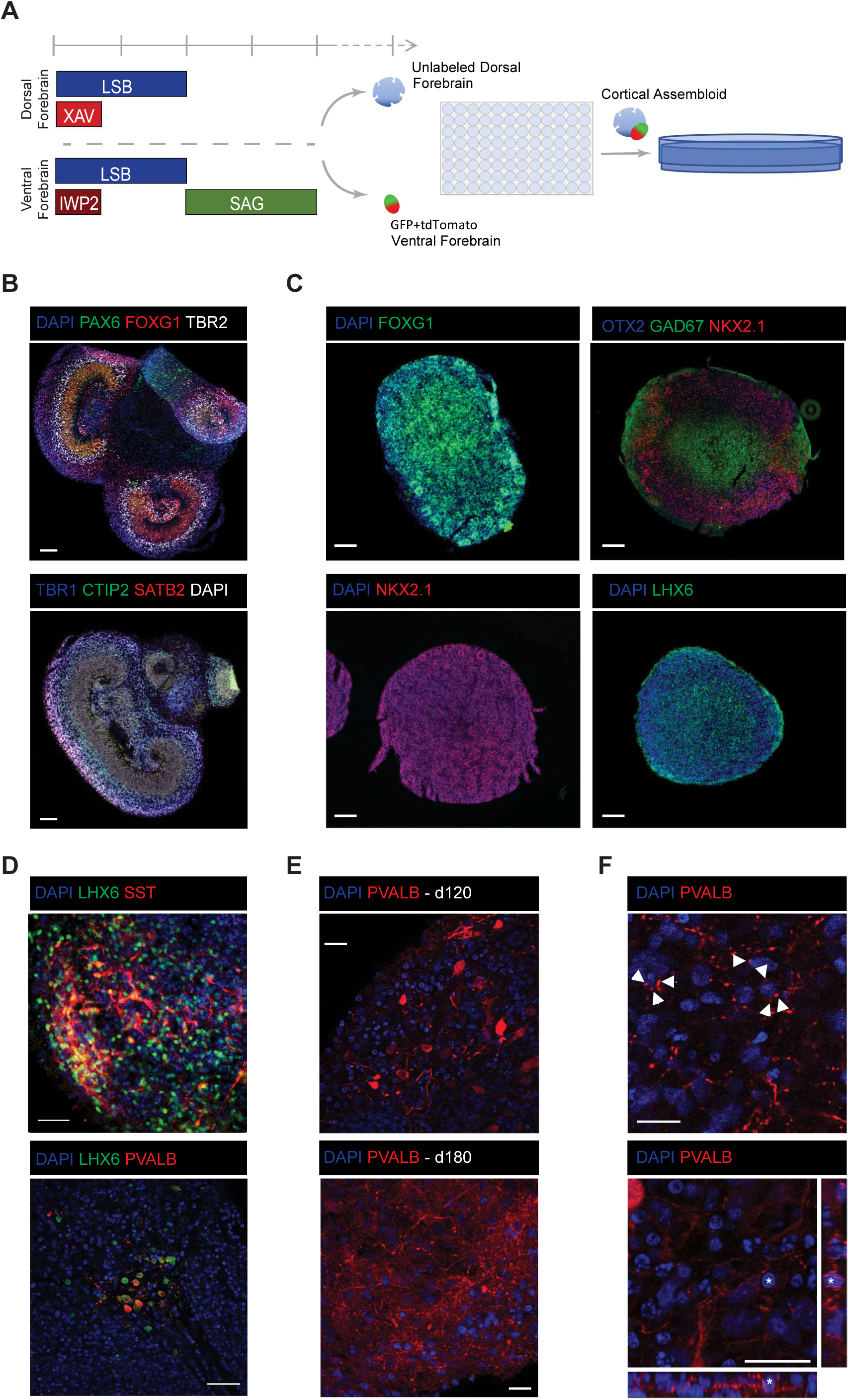
Patterning of dorsal and ventral forebrain organoids. **A)** Paradigm for patterning dorsal (top) and ventral (bottom) forebrain organoids. Organoids are transferred to a 96 well plate at d30 (middle) and then 10cm with rotation after 24hr (right). LSB: Dual SMAD inhibition; XAV:WNT inhibitor; IWP-2: WNT inhibitor (PORCN); SAG: Sonic Agonist. **B)** Dorsal forebrain organoid at d60 stained for patterning markers (left panel) and markers of cortical layers (right two panels). C) Ventral forebrain organoids at d16 ubiquitously express ventral forebrain markers (top and bottom left panels) and at d30 display expression of MGE-ventricular zone markers, NKX2.1, OTX2, physically separated from markers expressed in the mantle zone of the MGE, GAD67, LHX6 (top and bottom right). **D)** Co-expression of MGE-derived CIN marker LHX6 with SST at day 60 (top) and PVALB at day 120 (bottom). **E)** PVALB expressing neurons at day 120 (left) and day 180 (right). **F)** Basket like morphology and perisomatic boutons (arrowheads) in PVALB+ cortical interneurons at day 120.

We have previously derived cortical interneurons in monolayer culture by treating forebrain-patterned neural progenitors with a ventralizing SHH signal starting at the onset of telencephalic marker expression^56^. To adapt this differentiation paradigm for use in organoid culture, we applied the SHH agonist SAG following neural induction at day 8 of differentiation (Fig. 2A, bottom) resulting in ventral forebrain (vFB) organoids. On day 16, the time point of SAG withdrawal, vFB organoids display a near uniform expression of FOXG1, OTX2 and, importantly, NKX2.1 (Fig. 2C), which marks progenitors within the medial ganglionic eminence (MGE). Further, at day 30, we observed expression of GAD67 and the MGE-derived cortical interneuron marker LHX6 (Fig. 2C). These GAD67/LHX6+ cells appeared physically separated from the NKX2.1/OTX2+ progenitors, and were found predominantly in the center of the organoid, with the progenitor zone located in the exterior region, somewhat reminiscent of MGE organization with progenitors organized along the ventricular zone and postmitotic neurons migrating away into the mantle zone^26^.

Having successfully generated dFB and vFB patterned organoids, we next sought to aggregate them to generate an integrated cortical assembloid. To this end, we allowed our dFB and vFB organoids to develop separately until day 30, a timepoint when we observed the appearance of a significant postmitotic cortical interneuron population (see Fig. 2C), and then aggregated dFB and vFB organoids 1:1 in a low-attachment 96-well for a period of 24-48 hours (Fig. 2A). We next assessed whether our cortical assembloids would support the continued maturation and development of cortical interneurons by assessing them for markers of mature MGE-derived interneuron subtypes. Indeed, we observed robust expression of SST in LHX6+ neurons in day 60 cortical assembloids (Fig. 2D, top). No PVALB+ neurons were not observed at this time point, consistent with the highly protracted (postnatal to adolescent) timeline of PVALB+ interneuron development in vivo. Remarkably, however, when cortical organoids were allowed to mature to day 120, we were able to consistently derive PVALB expressing LHX6+ neurons (Fig. 2D, bottom). While PVALB expression has been observed in hPSC-derived cultures *in vitro* cultures previously^42–44,56–60,75^, convincing co-expression with MGE-derived cortical interneuron marker LHX6 has not been demonstrated. LHX6 co-expression is critical in confirming interneuron identity given expression of PVALB in other neuronal lineages (discussed above). Furthermore, in studies where there is evidence of potential PVALB+ cortical interneurons, they are only observed at much later stages of organoid development (> 6 months)^42,57,59^. In the current study, we show that it is possible to derive these neurons on a reduced time scale and that they continue to develop and display increasing complexity in later stage cortical assembloids (Fig. 2E). Further, we found evidence of perisomatic boutons (Fig. 2F; Suppl. Fig 1) starting in day 120 assembloids which are a hallmark of PVALB+ basket cells; a subtype of cortical interneuron believed to be important in mediating cognition^3^.

### Cortical assembloids support the development of cortical interneurons similar to those observed *in vivo*

In order to track vFB derived cells in an aggregated cortical assembloids, we generated isogenic hESC lines expressing either H2B:GFP or H2B:tdTomato using a P2A fusion in frame with the housekeeping gene GPI (Suppl. Fig. 2A&B). This strategy ubiquitously labeled hESCs and their progeny (Suppl. Fig. 2C&D), and reporter expression was maintained throughout the course of the differentiation even in the most mature neurons. We then generated vFB and dFB organoids alternately labeled with these ubiquitous reporters, aggregated them to form cortical assembloids (Fig. 2A), and could readily track the extensive migration of vFB-derived cells throughout the dFB. Interneuron progenitors often migrated from the vFB in large swaths, reminiscent of the migratory streams observed *in vivo*, and then spread out within cortical-like regions of excitatory neurons within the dFB organoid (Suppl. Video 1). Time-lapse imaging of cytoplasmic GPI-GFP labeled cortical interneurons migrating in cortical assembloids revealed morphologies characteristic of tangentially migrating cortical interneurons (Suppl. Video 2) – the extension and bifurcation of a branched leading process, a swelling in front of the nucleus in that process followed by saltatory nucleokinesis and retraction of the trailing process and one of the leading branches^76,77^. To further assess the diversity and development of cortical interneurons derived within this system, we performed single-cell RNA sequencing (scRNA-seq) on isolated vFB derived cells. At day 75, cortical assembloids generated with a GPI-H2B:GFP/tdTomato labeled vFB were dissected to separate the original vFB domain from the dFB domain and allow us to distinguish between vFB-derived cells that either migrated into the dFB portion of the assembloid or those that remained within the vFB portion. The separate vFB and dFB domains were dissociated and FACS was used to isolate fluorescently labeled cells for scRNA-seq. As expected, we observed multiple stages of MGE and cortical interneuron development, including an NKX2.1+/FOXG1+ population of MGE- progenitors, an intermediate stage of LHX6+/GAD1+/ERBB4+/DCX+ post-mitotic maturing cortical interneurons that lack subtype specific markers and a population of more mature SST+/LHX6+/GAD1+ cortical interneurons (Fig. 3A). Notably, the vFB-derived cells isolated from the dFB domain were primarily either immature post-mitotic or SST+ cortical interneurons whereas most of the MGE-progenitors were located in vFB domain (Fig. 3A). We identified additional, more minor cell populations that reflected LHX8+ striatal interneurons, which are also derived from MGE-progenitors, and neurons of the globus pallidus and striatum, all of which appeared strongly enriched in cells isolated from the vFB domain (Suppl. Fig. 3A&B). Finally, we observed another minor population representing CGE-derived cortical interneurons located in the dFB domain (Suppl. Fig. 3C). Together, these data indicate that our vFB organoids can produce the expected cell types of the ventral telencephalon, and that each of these cell types properly sorts between ventral and dorsal forebrain domains within the assembloid mimicking their corresponding *in vivo* developmental behavior.

**Figure 3.**
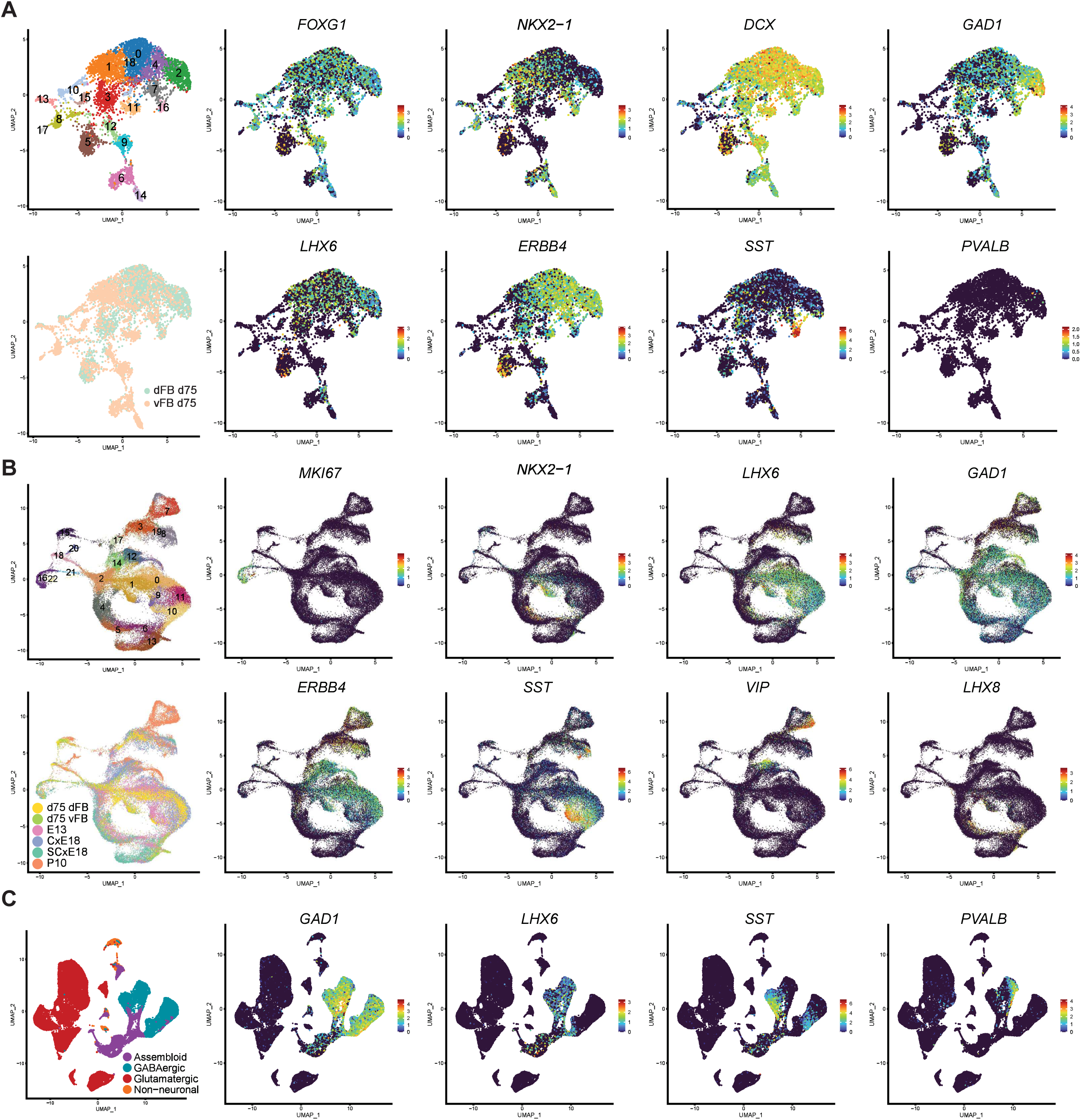
Molecular assessment and comparison of neocortical organoid derived CINs to their counterparts *in vivo.* **A)** Genes related to CIN development from scRNA-seq data of vFB-derived cells isolated from integrated neocortical organoids at day 75. **B)** scRNA-seq data integration with developing murine interneuron data from Mayer *et al.* 2018. C) Integration with adult postmortem motor cortex data from Bakken *et al.* 2021.

We next wanted to ask how closely the cell populations emerging within our cortical assembloids matched those of the developing and mature cortex *in vivo*. To this end, we first compared our combined dataset for dFB and vFB isolated cells against a previously published dataset from Mayer *et al*.^78^, comprising multiple stages of murine cortical interneuron development *in vivo*, using anchor-based data integration^79,80^. We observed significant overlap between multiple stages of cortical interneuron development in our cortical assembloids with their *in vivo* counterparts (Fig. 3B). Specifically, at day 75, the most mature LHX6+/GAD1+/SST+ cortical interneurons shared cluster 10 & 11 with murine SST+ cortical interneurons isolated from the dorsal forebrain from the Mayer et al dataset (Fig. 3B). Similarly, we observed postmitotic immature LHX6+/GAD1+/ERBB4+/SST- clustering together with in clusters 1 & 2 and MKI67+ progenitors clustering with subcortical progenitors of the Mayer et al dataset in cluster 16 (Fig. 3B). We also observed a cluster of VIP+/CCK+ CGE-like interneurons clustering with murine CGE interneurons in cluster 7 (Fig. 3B; Suppl. Fig. 3D), subcortical LHX8+ neurons sharing a cluster with striatal interneurons in clusters 5 and 6 (Suppl. Fig. 3D) and AQP4+ astrocytes in cluster 15. We next investigated how our assembloid-derived cortical interneurons compared to mature human cortical interneurons. We again performed anchor-based integration between the day 75 cortical assembloid data and published postmortem single nucleus RNA-seq data from the Allen Institute collected from multiple regions of the human cortex^81^. There was far less overlap with the cells from our dataset, which likely represents the temporal mismatch of fetal-like development in vitro versus the postmortem human data collected from adults. Nevertheless, we did observe a portion of our most mature cortical interneuron population clustering with either the SST+ or the PVALB+ cortical interneurons of the Allen Institute’s dataset (Fig. 3C). This was especially interesting since at day 75 of differentiation, the cortical interneurons in those assembloids have not yet begun expressing PVALB. Other populations where we observed significant overlap between the cortical assembloid dataset and the adult dataset from the Allen Institute represent astrocytes, oligodendrocyte precursor cells and a separate population of CGE-derived interneurons (Fig. 3C, right panels; Suppl. Fig. 3E). Overall, these data indicate that cortical interneurons developing within our cortical assembloids closely resemble those developing *in vivo*, and further, that the more mature cortical interneurons within our culture system segregate into populations of human SST+ versus PVALB+ cortical interneuron subtypes.

### Cortical assembloids show electrophysiological fast-spiking PVALB+ cortical interneurons structured network oscillations

The relatively early time point at which PVALB+ interneurons arose in our cortical assembloids afforded us a unique opportunity to assess the electrophysiologic properties of these cells. To achieve this, we generated a GFP-T2A-PVALB reporter by using CRISPR/Cas9 to knock in a GFP-T2A construct in frame with the 5’ end of the endogenous *PVALB* locus (Fig. 4A). We confirmed the reporter’s functionality in day 120 cortical assembloids generated by fusing WT dFB organoids to GFP-PVALB vFB organoids. We observed a 1:1 overlap of GFP and PVALB protein expression by IHC (Fig. 4B). We next used this reporter cell line to perform whole cell patch-clamp electrophysiology on GFP-PVALB+ cells in slices acutely cut from cortical assembloids (Fig. 4C-F). Notably, we observed firing patterns that are characteristic to fast-spiking PVALB+ cortical interneurons^82^, including delay to first spike (average delay 0.34 ± 0.12s, n = 6 out 12 cells; Fig. 4F), non-accommodating firing patterns, near-threshold ramping (average ΔV = 5.2 ± 0.6mV, n=12), near-threshold oscillations, stuttering firing patterns, and maximum firing frequencies of up to 78 Hz (Fig. 4E-G). While those frequencies are not as fast as expected for a fully mature fast-spiking PVALB+ cortical interneuron, they are beyond what would be expected from a non-fast-spiking interneuron in a cortical assembloid^42^.

**Figure 4.**
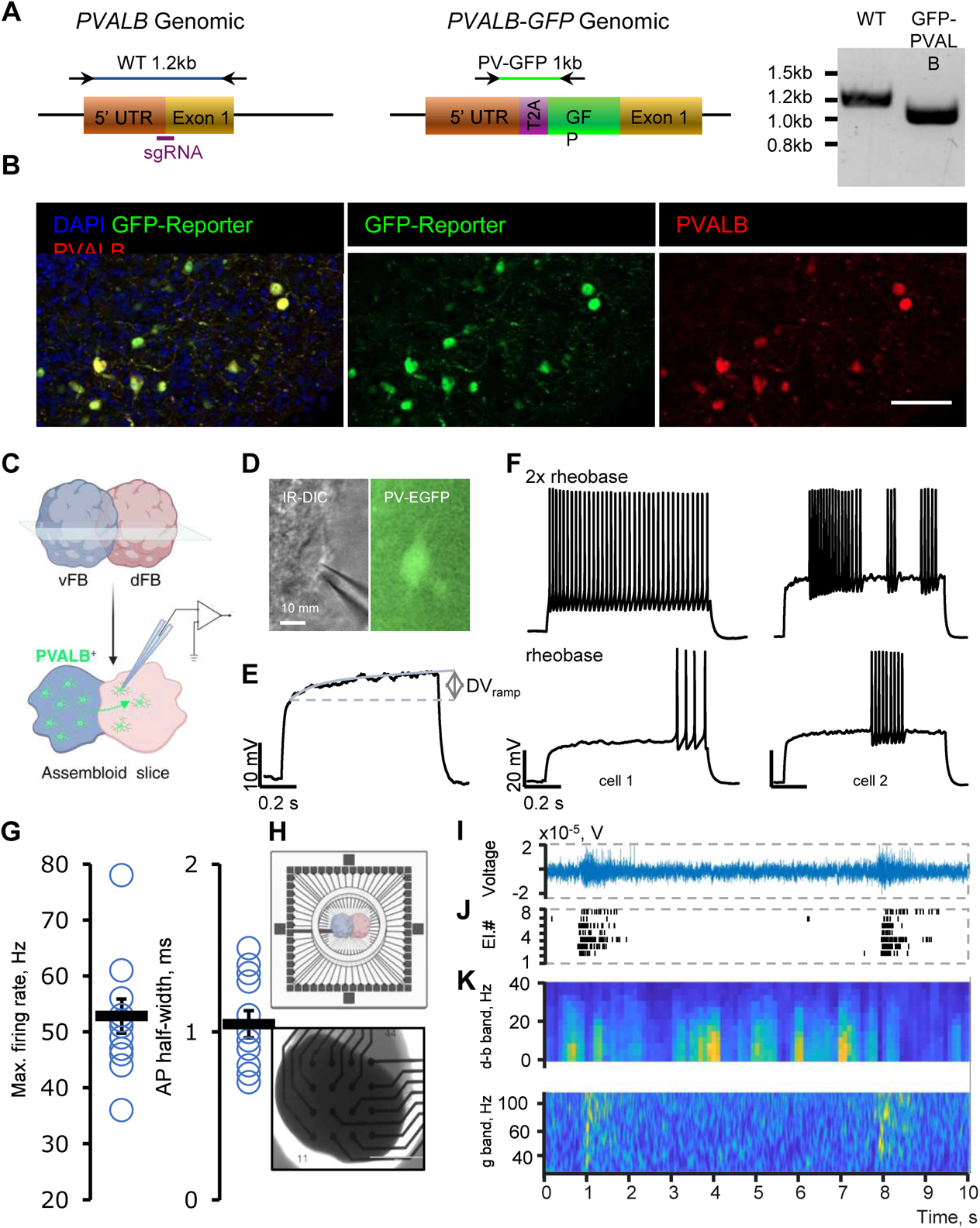
PVALB+ fast-spiking properties and γ-band oscillation in cortical assembloids. **A)** Schematic of strategy used to generate the PVALB reporter with expected genotyping bands displayed above. Genomic PCR for homozygous examples of each genotype shown to the right. **B)** Representative immunofluorescent imaging of the GFP version of the PVALB reporter in neocortical organoids at day 110 of differentiation. The organoid section shown was co-stained with GFP and PVALB to assess the reporter’s fidelity. Scale bar=S0um. C) Schematic of PVALB-GFP assembloids sliced and recorded by patch-clamp. **D)** Example of a clamped PVALB-GFP neuron in an assembloid slice. **E)** Near threshold oscillations in a PVALB+ neuron. **F)** Representative examples of characteristic spiking patterns observed in PVALB+ neurons. **G)** Quantification of max firing rate and action potential {AP) half-widths recorded from PVALB-GFP+ cells. **H)** Schematic of an assembloid recording on a muti-well MEA plate (top). Transmitted light image of a FB assembloid in a well of a 48-well MEA plate (bottom). Scale bar: 1000 µm. I) Electrical activity from a representative MEA electrode recorded during an organoid-wide network activity bursts. **J)** Raster plot of individual spikes sorted from all active electrodes in the well. **K)** A compound spike-time activity histogram from the electrodes shown in **J** that is overlaid onto the continuous wavelet transfer {CWT) of the recorded voltage waveforms. Color bar indicates CWT power (range 0-0.15).

To further evaluate cortical assembloid functionality at the network level, we performed recordings of the local field potentials (LFP) and spontaneous spiking activity using multi-electrode arrays (MEAs, Fig. 4H). Each cortical assembloid (n=8) was plated in a well of an MEA plate containing 16 low-impedance platinum microelectrodes. Recorded voltage signals (Fig. 4I) were differentially analyzed into neuronal firing (represented as a multi-unit activity) and the spectral power of the LFP signal which is separated between low-frequency (1-30 Hz, δ-β bands) and higher frequency bands (40-100 Hz, γ-band). After growing for 21 days on the MEA plate (total DIV = 128 days) forebrain assembloids displayed robust activity (mean firing rate 1.6 ± 0.9 Hz) that was organized into bursts of network-synchronized spiking interspersed with periods of quiescence (Fig. 4J). These network bursts were periodic (mean frequency 0.11± 0.03 Hz) with significantly higher intra-burst spiking activity levels (mean frequency 29 ± 8 Hz, Fig. 4J). LFP measurements in forebrain assembloids revealed simultaneous sustained oscillations at multiple frequencies ranging from 1-100 Hz (Fig. 4K). Interestingly lower frequency δ-β bands showed increased power during quiescent phase (Fig. 4K top), while γ-band oscillations were mostly present during network burst events (Fig. 4K bottom).

Our findings suggest that cortical assembloids are capable of generating PVALB+ cortical interneurons in as little as 120 days with relevant electrophysiological features and that that these assembloids have PVALB-relevant network oscillatory patterns.

### The 22q11.2 deletion impairs tangential migration of developing cortical interneurons

Our study presents a powerful modeling platform to study cortical interneuron development from the stages of early MGE patterning and tangential migration all the way to the acquisition of SST+ of PVALB+ fast spiking interneuron identities and function. Furthermore, we have generated a resource of isogenic hESC lines carrying schizophrenia-associated SVs. As an initial proof of concept for modeling potential schizophrenia-related disease phenotypes, we focused on three isogenic models: 22q11.2 del, 15q13.3 del, and 2q34 del. As an early functional readout, we assessed the potential impact of each of those SVs on disrupting tangential migration of cortical interneuron progenitors. We generated single vFB organoids containing equal mixes of isogenic WT and SV lines alternately targeted for tracking with either the GPI- H2B-GFP or GPI-H2B-tdTomato for cell tracking and generated cortical assembloids according to the paradigm displayed in Fig. 2A. We maintained labeled, chimeric WT:SV cortical assembloids to day 60 (30 days post-aggregation), then fixed and labeled them using the iDISCO tissue clearing protocol on intact assembloids^83^. Entire iDISCO-labeled assembloids were imaged via light sheet microscopy (Suppl. Video 3). For each SV, 3 independent clones were used to assess impact on migration, each experiment was run for 3-5 independent differentiation, and 3-5 assembloids were quantified per clone each time. To determine if an SV was associated with a defect in cortical interneuron tangential migration, we identified cortical-like regions of interest within the assembloid, which contains layers of excitatory neurons emanating from one to two large neural rosettes (Fig. 5A; also observable in Fig. 2B). We performed a paired analysis of the density of WT versus SV cells in cortical like regions of each assembloid and used the relative representation of each genotype within unaggregated vFB organoids from the same batch for normalization. Importantly, we observed a consistent and highly significant reduction in the normalized density of cells harboring the 22q11.2 deletion when compared to WT isogenic controls within the same assembloid (Fig. 5B) In contrast, both 15q13.3 del and 2q34 del SVs did not show any significant defects in migration.

**Figure 5.**
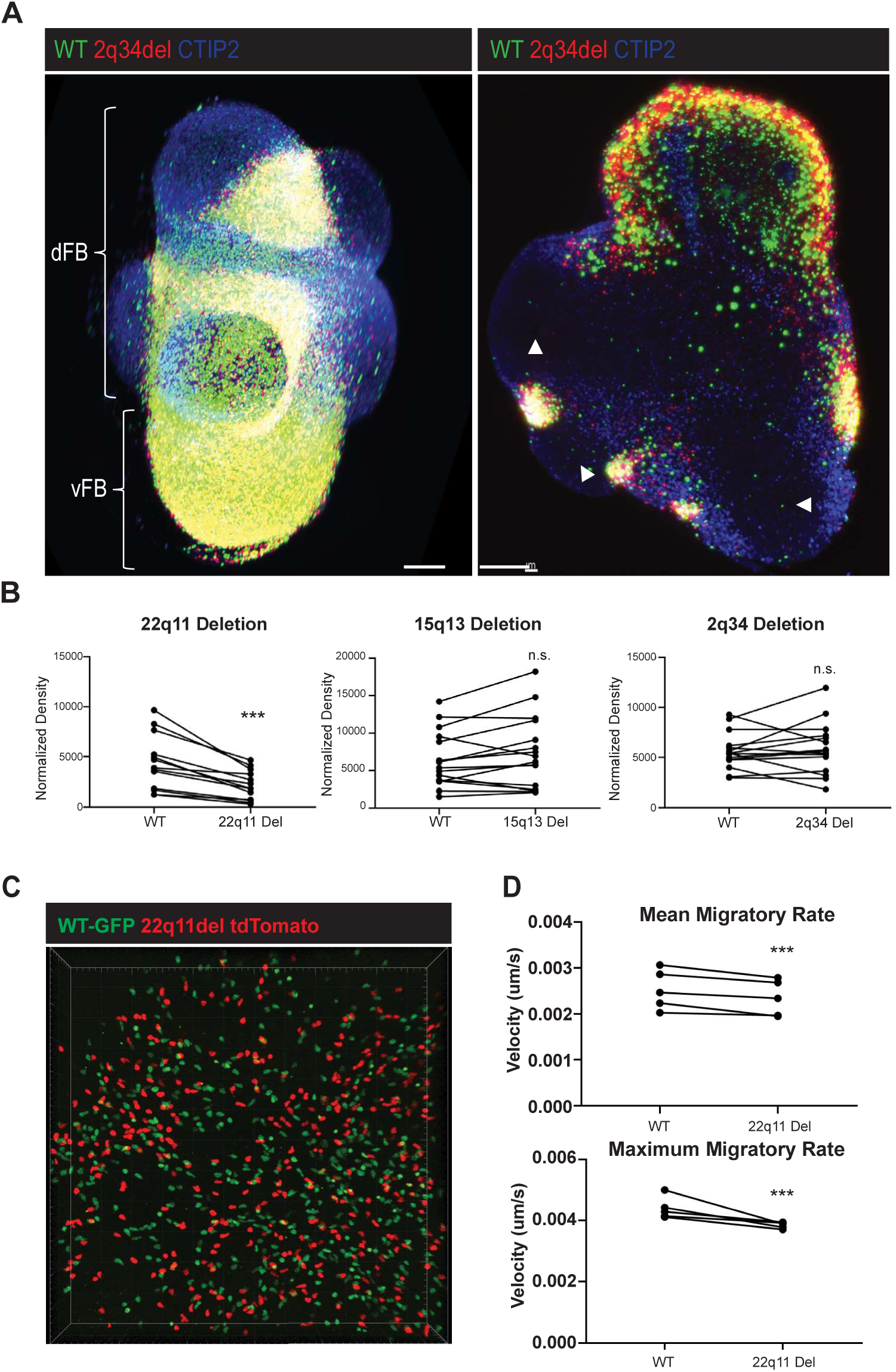
The 22q11.2 deletion leads to migratory defects in integrated neocortical organoids. **A)** Representative example of a day 60 3D iDISCO-cleared organoid imaged with the light sheet. (Left panel) Maximum intensity projection of cortical assembloid, dFB and vFB regions are indicated by brackets and “Cortical-like” regions which were used for quantification marked with *. (Right panel) Digital slice of a cleared light sheet imaged organoid, ventricular-like zones from which the cortical-like areas are generated are indicated with arrow heads. Scale bar=200um. **B)** Quantification of CIN density from cortical like areas at day 60. Each pair represents average normalized CIN density in cortical-like areas of a single organoid. C) Representative example of live image taken at 1 week post aggregation. Gridlines=S0um **D)** Quantification of migratory rates of CINs in the dFB region 1 week post aggregation, both mean (top) and maximum (bottom) migratory rates are shown. Each pair represents the average migratory rate of CINs within a single organoid. For the purposes of this analysis, immobile CINs, which may have already completed their migration or be yet to begin it, were excluded. Statistical significance for these data was determined with a paired student’s T-test; ** *= p<0.001; *=p<0.05.

To further characterize the tangential migration defect in the 22q11.2 deletion lines, we used confocal microscopy to perform time-lapse imaging on cortical interneurons migrating into the dFB portion of the cortical assembloids. Again, the GPI-H2B-GFP and GPI-H2B-tdTomato reporters were utilized to allow tracking of 22q11.2 deletion harboring cortical interneurons and their isogenic controls within the same assembloid (Fig. 5C). Consistent with our iDISCO observations, we observed a reduction in both the mean and maximum migratory rate of 22q11.2 deletion cortical interneurons when compared to isogenic controls (Fig. 5D, Suppl. Video 4).

Finally, we performed patch-clamp electrophysiology to assess for potential functional defects in 22q11.2 deletion cortical interneurons after migration. We noticed a progressive increase in the maximum number of action potentials fired by vFB neurons recorded between days 80 and 120, and that vFB-derived neurons in day 120 cortical assembloids could produce robust excitatory and inhibitory post-synaptic events (Suppl. Fig. 4A&B). We, therefore, selected day 120 for whole-cell patch-clamp recording of WT GPI-GFP and 22q11.2 deletion GPI-tdTomato labeled vFB-derived neurons within cortical-like dFB regions of the assembloids. Cortical assembloids from three distinct clones were recorded in three separate experiments; however, we observed no significant difference in the maximum number of action potentials generated between 22q11.2 deletion neurons and corresponding controls (Suppl. Fig. 4C). These data suggest, that the 22q11.2 deletion does not affect overall excitability of cortical interneurons at this stage of development; but given the migration deficit, fewer of them will reach their intended positions within the cortex, and thereby result in mis-formation of cortical micro-circuitry and alterations in cortical inhibitory control.

### Mitochondrial defects underly the tangential migration deficit in 22q11.2 deletion cortical interneurons

To explore the mechanisms behind the tangential migration defect in 22q11.2 del cortical interneurons, we again generated chimeric cortical assembloids with a vFB component comprising GFP- and tdTomato-labeled WT and isogenic 22q11.2 deletion lines. We next allowed these cortical assembloids to mature to day 60 of differentiation, the time point at which we observed an underrepresentation of 22q11.2 deletion neurons within the cortical-like areas of the assembloids, at which point they were dissociated and GFP+ and tdTomato+ cells were isolated via FACS for scRNA-sequencing. The 22q11.2 locus contains about 44 coding genes, plus additional noncoding transcripts. We could detect 40 of these in our dataset and found a significant global depletion for those genes in expression in the 22q11.2 deletion assembloids (adj. pval < 0.05; Suppl. Fig. 5A). To computationally identify a putative population of immature migrating interneurons within the single cell data, we performed RNA Velocity and cell latent time calculations (Fig. 6A), which accounts for splicing dynamics in determining a cell’s position along a trajectory. Trajectory analysis successfully identified a dividing *MKI67*+/*VIM*+ population of cells at its root and branching into 4 endpoints, one containing *AQP4*+/*VIM*+ astrocytes, another containing subcortical *LHX8*+ interneurons and two final branches representing cortical interneurons (Fig. 6A; Suppl Fig. 5C).We identified clusters within the cortical interneuron developmental trajectory with an intermediate latent time value (between 0.25 and 0.5; Suppl. Fig. 5B), at which cells were positive for post-mitotic cortical interneuron markers (e.g. *LHX6, GAD1* and *ERBB4*; Fig. 6A), but did not yet strongly express synaptic markers (e.g. *SYN1*, *GPHN* and *HOMER1*; Suppl. Fig. 5C) nor mature subtype markers (e.g. *SST* and *PVALB*; Fig. 6A; Fig. 6A; Suppl. Fig. 5C). We selected clusters 0 and 4 as the most representative for immature cortical interneurons, as they met the above criteria. We therefore reasoned that comparing 22q11.2 deletion cells and their isogenic controls within this specific cluster may reveal mechanistic insights into the tangential migration defect.

**Figure 6.**
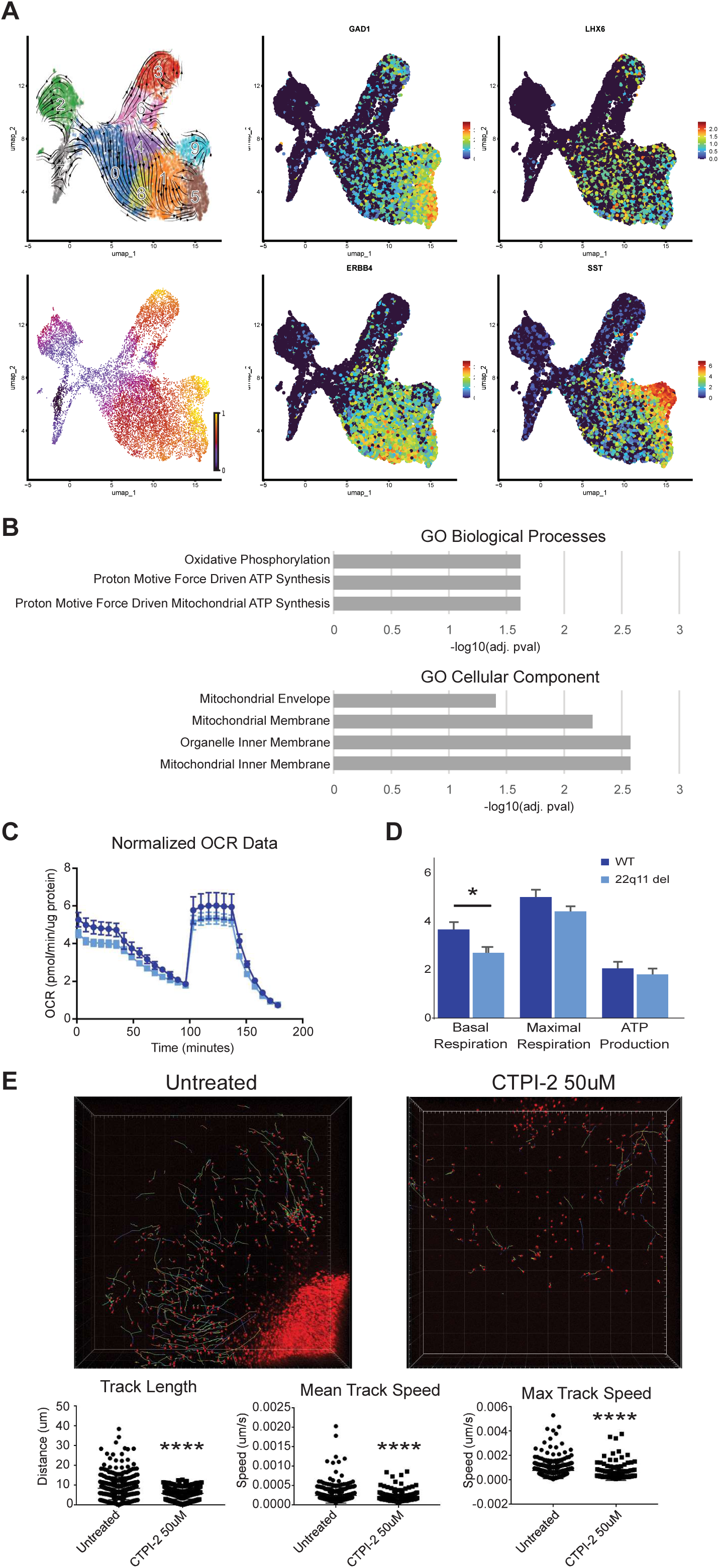
Migrating CINs harboring the 22q11.2 deletion exhibit mitochondrial defects. **A)** Velocity trajectory (top left) and latent time (bottom left) and marker expression (right 4 panels) analysis used to identify putative immature migrating populations of MGE-derived cortical interneurons. **B)** GO Biologic processes (top) and Cellular Compartment (bottom) terms downregulated in 22q11.2 deletion cells from cluster 4 compared to isogenic controls. No significantly enriched terms were observed for upregulated genes. C) Representative example of oxygen consumption rates (pmol/min) normalized by ug/protein from Seahorse organoid Mitostress assay normalized by ug of protein per organoid. **D)** Quantification of basal respiration, maximal respiration and ATP production from 3 independent Seahorse experiments. Each experiment recorded values from 5-8 organoids per condition. *= p<0.0SE) Traces from live imaging of CIN tangential migration in neocortical organoids treated with CTPl-2. Organoids were treated with at the onset of imaging, time-lapse images were acquired every 20min for 16hr. Tracks in the top panels trace the migration paths of WT cells over the time-lapse. The time-lapse was taken for 3 independent clones, shown are representative images and quantifications for a single clone. GPI-H2B:GFP=22del; GPI­ H2B:tdTomato=WT. grid lines= 100um.

While our study reveals a cortical interneuron tangential migration defect directly in human cells, animal models of the 22q11.2 deletion have previously suggested a cortical interneuron deficit^30,31^. One proposed mechanism in those mouse studies is the misregulation of CXCR4; specifically, that haploinsufficiency of the 22q11.2 gene DGCR8 leads to an indirect downregulation of CXCR4^30,31^. Surprisingly, however, we observed no evidence of CXCR4 downregulation in our 22q11.2 deletion immature cortical interneurons, nor did we note any reduction in any other signaling receptor/ligand interaction that has been reported as involved in cortical interneuron migration or homing (Suppl. Fig. 5D). When we expanded our analysis to include all genes differentially expressed between 22q11.2 deletion and their isogenic controls, we could detect multiple genes from the 22q11.2 locus downregulated within each of these clusters (*RANBP1, MRPL40, SLC25A1, DGCR6L* and *ZNF74* for cluster 0 and *RANBP1, MRPL40, SLC25A1, COMT* and *ZNF74* for cluster 4) as well as a general trend for those that did not reach statistical significance (Suppl. Fig. 5E). We performed GO analysis on the differentially expressed genes for each cluster, and we noted a consistent downregulation of terms related to mitochondria and mitochondrial function in cluster 4 (Fig. 6B), suggesting that haploinsufficiency of *SLC25A1* and *MRPL40* may be linked to a broader mitochondrial defect in these cells. While no GO terms reached statistical significance in cluster 0, which may represent a slightly earlier stage of interneuron development (Suppl. Fig. 5B), we were nonetheless able to observe a downregulation of *SLC25A1* and *MRPL40* in these cells.

We next determined whether the observed misregulation of mitochondrial genes did, in fact, lead to a functional defect. To this end, we measured OXPHOS in 22q11.2 deletion and isogenic control derived vFB organoids with an organoid-specific Seahorse Mito Stress Test. We noticed a defect in the OXPHOS pathway in 22q11.2 deletion vFB organoids, most notably a reduction in their basal metabolic rate (BMR; Fig. 6C&D). To further investigate the link between mitochondrial disruption and tangential migration, we employed the SLC25A1 inhibitor CTPI-2 in WT cortical assembloids during cortical interneuron migration. We first assessed toxicity of CTPI-2 treatment on assembloids and found no evidence of increased cell death during the treatment period and specific concentrations used (Suppl. Fig. 6A&B). Using our GPI- H2B:GFP reporter, we tracked migratory rates of vFB derived cells within the dFB portion of the cortical assembloid and, notably, observed a striking defect in CTPI-2 treated WT cortical interneuron migratory rates when compared to untreated controls (Fig. 6E). These data indicate that deficits in mitochondrial energy processing, likely related to *SLC25A1* and *MRPL40* haploinsufficiency, present a plausible mechanism for the 22q11.2 del-related cortical interneuron migration defect.

### The 15q13.3 deletion causes molecular defects specific to the PVALB+ subtype of cortical interneurons

Having identified a tangential migration defect for the 22q11.2 deletion during early cortical interneuron development, we next asked whether our cortical assembloid system could identify defects at later stages of development and focusing specifically on PVALB+ cortical interneurons. Mouse models of the 15q13.3 deletion have previously identified functional defects in PVALB+ cortical interneurons^84,85^. Therefore, we next performed scRNA-seq in cortical assembloids at day 125 for cortical interneurons carrying 15q13.3 deletion versus their isogenic controls, as described above for the 22q11.2 deletion at day 60 of differentiation. We sorted 15q13.3 deletion SV and isogenic control lines, alternately labeled by GPI- H2B:GFP or GPI-H2B:tdTomato, out of the same assembloids for sequencing. Similar cortical interneuron and glial populations were detected in the cortical assembloids at day 125 compared to day 75, with the notable addition of a sizable *PVALB+/LHX6+* population (Fig. 7A; Suppl. Fig. 7A-DA). We found that this *PVALB+/LHX6+* population expressed canonical cortical interneuron markers, similar to the *SST+/LHX6+* population. Within the *PVALB+/LHX6+* population we detected the upregulation of several potassium channels associated with PVALB+ interneuron firing properties in this population (Fig. 7B). Those include most notably *KCNC1* & *KCNC2* (Kv3.1, Kv3.2), which are required for sustained high-frequency firing^86^, *KCNA1* (Kv1.1), which is associated with spike-delay in delayed PVALB+ interneurons^87^, and *KCNK3* which has been shown to be involved in sub-threshold oscillations^82^. The expression of these channels correlated with the firing dynamics we previously observed for PVALB+ cells (Fig. 4C&D). Finally, within the PVALB+ compartment, we observed evidence for further substratification into potential cortical interneuron molecular subtypes, which included clusters expressing some markers previously associated with either basket or chandelier cell identity (Fig. 7C)^88,89^.

**Figure 7.**
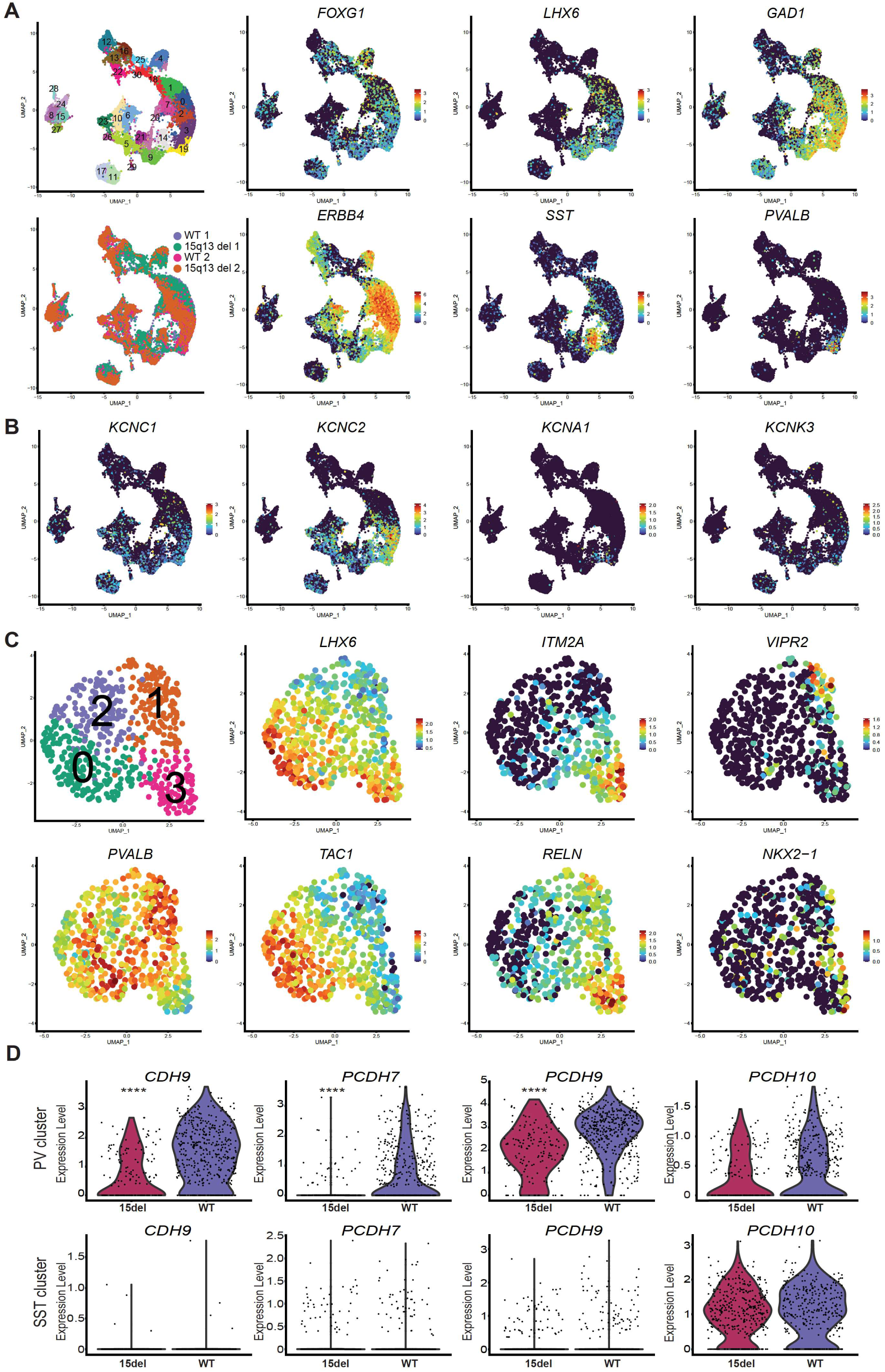
Molecular identity and defects of PVALB+ cortical interneurons harboring the 15q13 deletion. **A)** Genotype (top) and Louvain clusters (bottom) of day 125 scRNAseq of cortical assembloids. **B)** Markers of cortical interneurons in the day 125 cortical assembloids. C) Potassium channels important for fast­ spiking cortical interneuron function are enriched in the *PVALB+* cortical interneuron cluster. **D)** Cadherin and protocadherin genes differentially downregulated specifically in the 15q13 deletion PVALB+ cluster and their relative expression in the PVALB cluster (left) vs. the SST cluster (right) ***=p<0.001, ****=p<l.0e-10.

Having confirmed a molecular signature of the PVALB+ cortical interneuron phenotype in our cortical assembloids, we next assessed differentially expressed genes between 15q11 deletion and isogenic control cortical interneurons in both the *PVALB+* (cluster 6) and *SST+* (cluster 12). As expected, we observed a decrease in expression for each of the genes from the 15q13.3 locus in both populations (Suppl. Fig. 7E). Remarkably, while there were only few gene expression changes within the *SST+* cluster, 15q13.3 deletion harboring neurons within the *PVALB+* cluster showed a significant decrease in multiple cadherin and protocadherin genes compared to their isogenic controls (*CDH9, PCDH7, PCDH9,* and a trend for decreased expression of *PCDH10;* Fig. 7D). This reduction was not observed in the SST+ population, suggesting that these findings are specific to PVALB+ cortical interneurons carrying the 15q13.3 deletion. Those represent an especially interesting group of genes, as many of them have been linked to schizophrenia. For example, *CDH9* has been specifically associated with schizophrenia^90^ while *PCDH10* has been linked to both schizophrenia^91^ and, along with *PCDH9*, to autism spectrum disorders (ASD)^92,93^. Indeed, 15q13.3 deletion has also been associated with ASD, and potential cortical interneuron defects may be similarly relevant for ASD pathogenesis. Finally, *PCDH7* has been associated with modulating antipsychotic responses in schizophrenia patients^94^.

## Discussion

While defects in cortical interneurons have long been observed in schizophrenia patients, it has been difficult to study these defects in human model systems due to a range of factors, including the lack of candidate genes with strong effect, confounding genetic backgrounds and the absence of an in vitro platform that reliably captures the development of human PVALB+ fast-spiking cortical interneurons. While prior *in vitro* modeling work for schizophrenia in hPSC-derived cortical neurons has established an interesting set of candidate disease mechanisms^45–51,53,54^, it is crucial to ensure that those mechanism are confirmed in an experimental system with access to *bona fide* human cortical interneurons. GABAergic neurons of many differing types exist throughout the brain^95^ and, several of those share the expression of a subset of markers with cortical interneurons, including PVALB ^61–65^. While some previous studies have derived cortical interneurons from NKX2.1+ precursors^56,57^, whether or not those cells truly captured the properties of PVALB+ cortical interneurons remains unclear. Here we have developed a culture system that efficiently produces functional PVALB+/LHX6+ PVALB+ cortical interneurons within 120 days of differentiation, and we have used this platform to study early cortical interneuron defects in isogenic SVs relevant to schizophrenia.

The exact reasons we were able to rapidly derive LHX6+/PVALB+ fast-spiking cortical interneurons in our cortical assembloids as compared to other culture systems remain to be determined. It is likely that multiple factors contribute to this difference, perhaps beginning with proper developmental specification of the MGE and the cortical environment for cortical interneurons to develop in. After the guided differentiation of the organoids, our cortical assembloids are maintained without additional growth factors and without Matrigel. In culture systems that lack PVALB+ interneurons or where PVALB expression is delayed, perhaps extended mitogen exposure (e.g., EGF & FGF2) causes delayed neuronal maturation. Other differences in base media composition or supplements between protocols could also impact maturation rates. While beyond the scope of the present study, in future studies, it will be interesting to determine the primary culprit for accelerated PVALB+ cortical interneuron production and maturation and to further enhance PVALB+ interneuron yield.

An imbalance in the E/I ratio in schizophrenia has frequently been attributed to dysfunction rather than loss of cortical interneurons. Our findings that the 22q11.2 deletion leads to a cortical interneuron migration defect and reduced representation of cortical interneurons within the cortical-like regions of the cortical assembloid suggests a potential alternate mechanism affecting the E/I ratio in 22q11.2 patients. Indeed, this observation is supported by a mouse model of the 22q11.2 deletion, which shows a similar cortical interneuron tangential migration defect^30,31^. Furthermore, while several studies which examined cortical interneurons in postmortem human tissues have reported a loss of cortical interneuron gene expression without a reduction in cortical interneuron number^10–13^, other studies have reported such a reduction^36–41^. Moreover, postmortem studies in 22q11.2 patients reported an increase in the incidence of periventricular nodular heterotopia (PNH)^96–98^, a disorder that results from dysfunctional or arrested neuronal migration. Notably, PNH in these patients appears to stem from dysfunctional cortical interneuron migration^96^. However, whether there is reduced cortical interneuron density in the cortex of these patients remains unclear. Moreover, an increase in GABAergic interstitial white matter neurons (IWMNs) has been observed in several postmortem studies of schizophrenia patients^99–104^. While the cause of increased IWMNs is not known, one possible mechanism could be faulty cortical interneuron migration during development. In support of this, one study demonstrated that the same group of patients with an increase in GAD65/67+ IWMNs showed a reciprocal reduction in either GAD65/67^104^ or SST^103^ RNA levels in the overlying cortical layers. Therefore, for a subset of schizophrenia patients, such as those with a 22q11.2 deletion, a cortical E/I imbalance can be the result of dysfunctional cortical interneuron migration. However, it will be important to determine for what fraction of schizophrenia patients this may be the case and to ask whether environmental or genetic factors during early development may lead to similar migratory defects. Our hPSC-based platform could present a powerful tool to address those points in the future.

The 22q11.2 locus contains genes for the mitochondrial ribosome, *MRPL40*, and the sole mitochondrial citrate transporter in humans, *SLC25A1*, both of which are important regulators of mitochondrial function^46,105^. The locus further contains genes coding for *PRODH*, *TXNRD2*, *T10* and *ZDHHC8* that were only lowly expressed in our scRNA-seq analysis, but also encode for proteins localized within the mitochondria^106^. There is evidence for reduced mitochondrial function in 22q11.2 deletion neurons, as previous reports demonstrated a reduction in mitochondrial gene expression and OXPHOS rates in excitatory cortical neurons^46,47,107^. Furthermore, mitochondrial function in 22q11.2 deletion neurons is inversely correlated with schizophrenia penetrance^47^. However, whether and how these defects may also disrupt cortical interneuron development was unknown. We demonstrate for the first time in a human experimental system that 22q11.2 hemizygosity leads to cortical interneuron tangential migration defects, and that those are linked to mitochondrial deficits in these cells. Interestingly, cortical interneurons appear to switch from relying on glycolysis to OXPHOS earlier in their developmental progression than excitatory neurons, and disruption of OXPHOS has been shown to impede their tangential migration, but not the radial migration of excitatory neurons^108^. Nevertheless, our study does not rule out that 22q11.2 hemizygosity leads to additional defects at later stages of cortical interneuron development which will be the focus of future studies.

Finally, we have identified misregulated expression of multiple cadherins and protocadherins specifically within the PVALB+ population of 15q13.3 deletion cortical interneurons. The closely related SST+ population of cortical interneurons displayed no such defect, highlighting the importance of capturing the proper disease-relevant cell type in our culture system. In addition to the published reports linking select members of this group with schizophrenia, cadherins and protocadherins play an important role in broad aspects of neural development including synaptogenesis and dendritic spine morphogenesis^109,110^. Prior models of the 15q13.3 deletion have shown reduced and abnormal dendritic spines in murine cortical neurons and human iNeurons^111–113^, though it is not clear from those studies whether cortical interneurons are affected by those or other disease-related phenotypes. It will be interesting to study whether 15q13.3 deletion PVALB+ cortical interneurons display potential dendritic spine defects and to explore the roles cadherins and protocadherins play in this process. However, bona fide spines are rarely reported in hPSC-derived neurons and either require extensive culture (>8months)^114^ or the introduction of exogenous factors^115^. Therefore, further technical advances will be needed to properly assess this phenotype.

In conclusion, we have developed a cortical assembloid system that mimics early forebrain development and supports the functional maturation of PVALB+ fast-spiking cortical interneurons in just four months This culture system presents a powerful tool for modeling schizophrenia, but also highly relevant for modeling epilepsy and ASD, where PVALB+ cortical interneurons are also strongly implicated. We further demonstrate in isogenic lines the impact of SV-specific defects on immature and *PVALB+* cortical interneurons, illustrating the utility of this platform in assessing disease-related effects at multiple stages of cortical interneuron development. While these defects were observed for specific schizophrenia associated deletions, they may apply to a broader subset of schizophrenia patients. The application of this approach in future studies should allow for the identification of patients with similar cortical interneuron defects and the identification of additional genetic and environmental factors driving schizophrenia risk.

## Acknowledgements

We would like to thank all the Studer lab members for their insightful comments and input on the work presented here. We would also like to thank the Integrated Genomics Operation (IGO), Molecular Cytology Core and Flow Cytometry Core at Memorial Sloan Kettering Cancer Center (MSKCC) and the Bio-Imaging Resource Center (BIRC) at Rockefeller University. This work was supported in part through NIH grants R01AG054720 and R01NS128087 and through support from the Starr foundation to L.S. R.M.W. was supported for this work by the NIMH F32 fellowship 5F32MH116590. G.W.C. was supported for this work by the NIMH grant R01MH097879 to J.A.G. We thank Dana Pe’er, Ronan Chaligné and Single Cell Analytics Innovation Lab (SAIL) at MSKCC for advice on single cell analysis. We further thank Gustav Cederquist, Jason Tchieu and Johannes Jungverdorben for advice on organoid technology and PSC differentiation approaches.

## Contributions

L.S and R.M.W. conceived and designed experiments, analyzed and interpreted data, and wrote the manuscript.

R.M.W. performed organoid/assembloid differentiations, generated SV and transgenic lines, performed fixed and live imaging experiments, iDISCO experiments, single cell analysis, and Seahorse experiments.

I.K. performed the PVALB electrophysiology and MEA experiments and analyses.

K.K. performed MEA experiments.

G.W.C. & J.A.G. performed and analyzed the 22q11.2 electrophysiology experiments.

L.J, S.Y.C. & G.C. performed some imaging experiments and analysis.

All authors provided feedback in editing the manuscript.

## Methods

### hESC culture

MEL1 hESC lines were maintained feeder-free on vitronectin (Thermo #A14700) coated plates in Essential 8 Flex media (Thermo #A2858501). hESCs were passaged as clumps with a 0.4% EDTA/PBS dissociation solution. Cells were cultured at 37°C at 5% CO_2_ with routine testing for mycoplasma. Genomic integrity was assessed with karyotyping and shallow whole genome sequencing.

### Transfection of hESC

hESCs were dissociated to a single cell suspension with Accutase (Innovative Cell Technologies # AT104) and plated at a density of 250,000cells/well in a vitronectin coated 6-well plate in Essential 8 Flex supplemented with 10 µM Y-27632 ROCK inhibitor (Bio-Techne #1254/50). The following day, media was replaced with 2ml mTeSR (STEMCELL Technologies #85850) supplemented with CloneR (STEMCELL Technologies #05888). Lipofectamine-DNA complexes were assembled in Opti-MEM (Thermo #31985062) according to the manufacturers protocol, using 8ul of Lipofectamine Stem (Thermo #STEM00001) and 5ug plasmid DNA per well. 24 hours post transfection, media was replaced with Essential 8 (Thermo #A1517001) supplemented with CloneR. 48 hours following transfection, hESCs were dissociated to single cells with Accutase and GFP+ cells that received the pX458 vector were isolated via FACS with a BD Aria6. Sorted cells were replated in vitronectin in Essential 8 with CloneR. CloneR was removed after 4 days according to the manufacturer’s protocol and clones were picked and assessed for either deletion or transgene incorporation.

### Generation of Structural Variants and reporter lines

The 22q11.2 and 15q13.3 deletions were generated by targeting flanking segmental duplications with a single sgRNA^116^. An sgRNA predicted to cut once within each of two flanking segmental duplications was selected (22q11.2 sgRNA: GTAGAAAGGGCTTTGACACG; 15q13.3 sgRNA: GGCTGACTACATCCAGGAAC) and cloned into the pX458 Cas9/sgRNA expression vector^117^. The 2q34 deletion was generated by targeting sequences flanking the patient deletion with two distinct sgRNAs (2q34 sgRNA 1: GAGTTTGCAGACAGAGTATA; 2q34 sgRNA 2: GCCTTGCATAATATTGAAAC). *GPI* and *PVALB* targeting constructs were generated from a genomic DNA PCR with Q5 polymerase (New England Biolabs #M0494) amplifying 400-600bp of homology per side and assembled with NEBuilder (New England Biolabs #E2621S). An sgRNA targeting *GPI* (GPI sgRNA: CTTCATCAAGCAGCAGCGCG) and *PVALB* (PVALB sgRNA: GACAGACTTGCTGAACGCTG) were co-transfected with their respective targeting constructs for line generation. hESC lines were transfected as described above, for reporter lines, a ratio of 1:5 of sgRNA vector to targeting vector was used. TaqMan Genotyping Master Mix (Thermo #4371355) with CNV assays (Thermo) targeting loci within and flanking each CNV were used to screen clones. Positive hits were further assessed with shallow whole genome sequencing (sWGS) to confirm the correct SV.

### Organoid Differentiation and assembly

hESCs cultures at 50-70% confluency were dissociated with either 6 minutes of EDTA dissociation solution at room temperature or 4 minutes Accutase at 37°C if single cells were desired. For differentiation, 10,000 cells were plated into V-Bottom low attachment dishes (S-Bio #MS-9096VZ) in Essential 8 with ROCK inhibitor and 5uM XAV939 WNT inhibitor (R&D #3748). At 18-24hours after plating, organoids were washed 1x with PBS and treated with 100nM LDN193189 (Stemgent #04-0074) and 10uM SB431542 (R&D #1614) and either 5uM XAV939 for dFB or 4uM IWP-2 (Stemgent #04-0034) for vFB in Essential 6 Media (Thermo #A1516401). XAV939 treatment was discontinued after 72hours and IWP-2 treatment after 24hours, Essential 6 LDN193189 and SB431542 treatment was continued for both conditions until day 8. On day 8, dFB organoids were transferred to Organoid Media - 1:1 mixture of DMEM/F12 (Thermo #11320033) and Neurobasal (Thermo #21103049) supplemented with 1X GlutaMAX (Thermo #35050079), 0.5X NEAA (Thermo #11140050), 1X Pen/Strep (Thermo #10378016), 55uM 2-mercaptoethanol (Thermo #21985023), 1ug/ml human insulin (Thermo #12585014), 0.5X CTS N2 supplement (Thermo #A1370701), and 0.5X MACS NeuroBrew-21 (Miltenyi #130-093-566) - while vFB organoids were switched to a ventralization media - DMEM/F12 supplemented with 1uM SB431542, 1uM SAG (Tocris #4366), 1X CTS N2 supplement, 1X Pen/Strep, 1X GlutaMAX, 1X NEAA). Ventralization media was maintained until day 16 of the differentiation at which point vFB organoids were switched to Organoid Media and both conditions were transferred from 96well to 10cm plates and placed on an orbital shaker set to 70-75rpm. On day 30 of differentiation, a single dFB was placed in a well of a low attachment V-bottom 96 well plate with a single vFB organoid to generate a cortical assembloid. After 24 hours, cortical assembloids were carefully transferred back to 10cm plates using a cut-tip P1000 pipette, this plate was kept without rotation for an additional 4 days, after which it was returned to the orbital shaker where cortical assembloids were cultured in Organoid Media indefinitely.

### Immunofluorescent labeling & imaging

Organoids were fixed for staining overnight in 4% paraformaldehyde 1X PBS at 4^°^C with rotation and then cryoprotected in 30% sucrose 1X PBS for > 24hours at 4°C. Cryoprotected organoids were snap frozen in OCT (Fisher #NC9806257) on dry ice and 30-45micro floating sections were taken on a Leica Cryostat. Stains were performed in suspension in 1.7ml tubes, sections were washed 10min in 1X PBS 0.3% Triton-X100, blocked in 1X PBS, 0.3% triton and 10% Normal Goat or Normal Donkey serum for 2hr. Primary stains were performed overnight at 4°C with rotation in 1X PBS 0.3% Triton-X100. Sections were washed at room temperature with rotation 3X in wash buffer (1XPBS 0.1% Tween-20), stained with secondary antibodies 2hr in wash buffer, then washed 3 additional times with DAPI added to the second wash prior to mounting on slides. Primary antibodies used: PAX6 (BD #561462; 1:500), FOXG1 (Takara #M227; 1:500), TBR2 (eBioscience # 14-4877-82, 1:200), CTIP2 (Abcam ab18465; 1:500), TBR1 (Abcam #ab183032; 1:500), SATB2 (Abcam #ab51502; 1:50), MEF2C (Cell Signal #5030; 1:400), GAD67 (Millipore # MAB5406; 1:1000), OTX2 (Neuromics #GT150595; 1:100), NKX2.1 (Abcam #ab76013; 1:500), LHX6 (SCBT #sc-271433; 1:150), SST (Millipore #MAB354; 1:100), PVALB (Swant #PV 27; 1:5000), SYN1 (Synaptic Systems # 106 104; 1:500), GPHN (Synaptic Systems #147 011), GFP (Abcam #ab13970; 1:2000), mCherry (Rockland #600-401-P16; 1:500), and mCherry (Thermo #M11217; 1:2000). Slides were imaged with either a Zeiss Axio Observer or Leica SP8 confocal microscope.

### iDISCO and light sheet microscopy

Fixed assembloids were stained with the iDISCO+ protocol with previously validated antibodies^83^. Briefly, fixed assembloids were permeabilized using the described alternative pretreatment that does not necessitate methanol treatment prior to antibody staining. Assembloids were treated with permeabilization solution, blocking solution, and primary and secondary antibody solutions for 24 hours each. Then embedded in 1% Agarose prior to dehydration and clearing with dichloromethane (Sigma #270997) and benzyl ether (Sigma #108014). Cleared assembloids were imaged in a benzyl ether filled chamber on a LaVision Ultramicroscope II.

### Live imaging

For live imaging experiments, cortical assembloids one week post assembly were immobilized to 35mm glass bottom dish (MatTek #P35G-1.5-14-C) by covering them with a drop of phenol red free growth factor reduced Matrigel (Corning #356231) and allowing it to solidify for 10min at 37°C. Z-stacks of images between 40 and 60um were collected on a Leica SP8 confocal microscope every 20 minutes for 14 hours in a humidified chamber at 37°C with 5% CO_2_ in phenol red free organoid media.

### Quantification of migrating interneurons in cortical assembloids

Imaris v8.4.2 was used for quantification of both cleared and live assembloids. For cleared assembloids, image stacks from day 60 assembloids were imported into Imaris to render a 3D object. The surfaces function was used to define regions of interest around cortical-like areas of neurons deriving from 1-2 rosettes within the dFB territory of the cortical assembloid. The spots function was used to independently identify vFB derived cells within the ROI for each channel. The raw counts for each channel divided by the volume of each ROI were taken for each assembloid. For normalization, 30-micron sections taken on day 30 (time 0 for assembly) from the vFB organoid batch used for assembloid construction, stained for tdTomato and GFP and imaged. Relative amounts of each genotype within assembloids from each batch were determined by quantifying GFP and tdTomato with Ilastik v1.4.0^118^ using the Cell Density Counting function. Raw densities per assembloid were normalized by multiplying by 1 minus the ratio of that genotype’s representation within the day 30 organoid.

For time lapse experiments, regions of interest were selected that contained both GFP and tdTomato cells but were within the dFB (unlabeled) portion of the cortical assembloid. Imaris’s spots function was used to identify and track cells of each genotype over time. Track length for cells that remained within the imaging plane, mean and max speed were all quantified. Cells that remained immobile for the entire time course were excluded from the analysis, as these were unlikely to represent migrating cortical interneurons.

### Single Cell RNA Sequencing

Cortical assembloids were washed with 1X PBS and dissociated at 37°C 5% CO_2_ on an orbital shaker with Papain (Worthington #LK003150) for 30 minutes, broken up by repetitive pipetting with a 5ml pipette and returning to the shaker for an additional 15 minutes and again further dissociated with repetitive pipetting. Papain was deactivated and washed using the manufacturer’s protocol. For the separate vFB and dFB sequencing experiments in Figure 3, assembloids were manually dissected prior to dissociation using a scalpel and a fluorescent scope to identify the original vFB and dFB regions of the assembloid. Dissociated cells were resuspended in 0.4% BSA 1X PBS with 0.2U/ml RiboLock RNAse Inhibitor (Thermo #EO0382) and fluorescent cells were isolated via FACS on a BD Aria6 using DAPI to exclude dead cells. Prior to sort, cells were diluted to 1,000 cells/ml in 0.04% BSA 1X PBS 0.2U/ml RiboLock. Single cell suspensions were stained with Trypan blue and Countess II Automated Cell Counter (Thermo) was used to assess both cell number and viability. Following QC, the single cell suspension was loaded onto Chromium Next GEM Chip G (10X Genomics #1000120) and GEM generation, cDNA synthesis, cDNA amplification, and library preparation of 7,000-15,000 cells proceeded using the Chromium Next GEM Single Cell 3’ Kit v3.1 (10X Genomics #1000268) according to the manufacturer’s protocol. cDNA amplification included 11-12 cycles and 30-240 ng of the material were used to prepare sequencing libraries with 10-16 cycles of PCR. Indexed libraries were pooled equimolar and sequenced on a NovaSeq 6000 in a PE28/88 run using the NovaSeq 6000 S1 or S4 Reagent Kit (100 or 200 cycles) (Illumina).

### Single Cell Data analysis

Demultiplexing and alignment to the GRCh38 reference genome was performed with the CellRanger pipeline (Version 6.1.2). Data analysis was performed with R v4.1 using Seurat v4.2.0^119^. Cells expressing between 200 and 4,000 genes and less than 15% counts in mitochondrial genes were used in downstream analysis. Gene counts were normalized by total counts per cell and ScaleData was used to regress out cell cycle gene expression variance as determined by the CellCycleScoring function. PCA was performed on scaled data for the top 5000 highly variable genes and a JackStraw significance test and ElbowPlot were used to determine the number of PCs for use in downstream analysis. A Uniform Manifold Approximation and Projection (UMAP) on the top 15 PCs was used for dimensional reduction and data visualization. FindNeighbors on the top 15 PCs and FindClusters with a resolution of 2.0 were used to identify clusters. For trajectory and pseudotime analyses, Velocity v0.17.17^120^ was used to define spliced vs unspliced reads, and loompy.combine (Loompy v2.0.17; http://loompy.org) was used to merge samples into a single loom file. Downstream analysis was carried out in Python v3.9.12 with Scanpy v1.9.1^121^ and scVelo v0.2.4^122^ scvelo.pp.filter_and_normalize(min_shared_counts =20, n_top_genes =3000) was used for normalization, and log transformation and moments and nearest neighbors were then calculated with 17cVelo.pp.moments, and batch correction performed with bbknn. Louvain clustering was performed and clusters displaying low library complexity or signs of ER/glycolytic stress^123,124^. Velocities were estimated using the stochastic model with 17cVelo.tl.veloctiy(mode = “stochastic”) and latent times were obtained from 17cVelo.tl.recover_dynamics and 17cVelo.tl.latent_time using the default velocity identified root cells as the root. Clusters for differential expression analysis were selected based on immature cortical interneuron marker expression and velocity determined latent time. Differential expression analysis performed with MAST within clusters in Seurat with FindMarkers with ribosomal genes filtered out. Enrichr ^125,126^ was used to identify GO Biological Processes and GO Cellular Component terms associated with genes that were significantly differentially expressed (adjusted p-value < 0.05).

### Electrophysiology

For electrophysiology experiments on PVALB+ interneurons, day 120+ assembloids were embedded in 1% agarose and a sliced into 250µm-thick sections in ice-cold dissection solution using a vibratome (Vibratome Inc, Series 1500). Dissection solution consisted of (in mM): 195 Sucrose, 10 NaCl, 2 NaH2PO4, 5 KCl, 10 glucose, 25 NaHCO3, 4 Mg2SO4, and 0.5 CaCl2, adjusted to pH 7.3. After dissection and prior to recording, slices were kept in dissection solution bubbled with the mixture of 95% O_2_ and 5%CO_2_. For whole-cell recordings, slices from cortical assembloids were visualized using an upright Olympus BX51WI epi-fluorescence microscope equipped with a water-immersion 40x objective and differential interference contrast optics. Slices were constantly perfused with BrainPhys® medium (STEMCELL Technologies, Catalog #05791) preheated to 30-31°C and oxygenated with the mixture of 95% O_2_ and 5%CO_2_. Patch electrodes (4-6 MOhm tip resistance) were filled with internal solutions containing (in mM): 130 K+ gluconate, 6 KCl, 4 NaCl, 10 Na^+^-HEPES, 0.2 K+EGTA; 0.3 GTP, 2 Mg^2+^-ATP, 0.2 cAMP, 10 D-glucose. The pH and osmolarity of the internal solution were adjusted to resemble physiological conditions (pH 7.3, 290­300 mOsmol). Current- and voltage-clamp recordings were carried out using a Multiclamp 700B amplifier (Molecular Devices), digitized with DigiData 1440A digitizer and fed to pClamp 10.0 software package (Molecular Devices). In cases where spontaneous EPS/IPS-currents were recorded, individual neurons were held at Cl^-^ or Na^+^ reversal potential of −75 or 0 mV correspondingly. Data processing and analysis were performed using ClampFit 10.0 (Molecular Devices) and MatLab (MathWorks) software. P-values were calculated using a two-tailed, unpaired t test.

For the MEA recordings, a 48-well MEA plate (Axion Biosystems) was coated with 100 µg/ml poly-L- ornithine and 10 µg/ml laminin. Assembloids were plated on day 107 at one per well of the MEA plate into 8 wells. The cultures were fed twice a week and measurements were collected 8 hours after the medium was changed, 3 weeks after plating (on day *in vitro* 128). Recordings were performed using a Maestro MEA system (Axion Biosystems) with bandpass acquisition set at (0..5000 Hz), digitized at 12.5 KHz and acquired with AxIS software. Multi-unit spikes were detected with AxIS software routine using an adaptive threshold crossing that was set at 5.5 times the standard deviation of the estimated noise for each electrode. For the MEA activity analysis, the electrodes that detected at least 5 spikes/min were classified as active electrodes. Network bursts were defined as events with minimum number of 5 spikes and inter-spike interval <100 ms occurring in at least 25% of active electrodes. LFP signals were lowpass filtered (Butterworth, <500Hz) and analyzed using custom MATLAB routines. To analyze lower band (1-30) LFP frequencies we used the continuous multitaper spectrogram function ‘mtspecgramc’ from the Chronux package (chronux.org). For the analysis of higher band (30 - 100Hz) LFP power spectrum we utilized continuous wavelet transform (cwt) function using Morse wavelet.

### Assessment of mitochondrial function

Mitochondrial function was assessed with the Seahorse XF Cell Mito Stress Kit (Agilent #103015-100) and analyzed on a Seahorse XFe96 (Agilent) according to the manufacturer’s protocol with the following modifications. Briefly, one vFB organoid generated entirely from either the 22q11.2 deletion line or isogenic control was placed in each well of a Seahorse XFe96 Spheroid Microplate (Agilent #102978-100) coated with poly-D-lysine (Sigma #P6407). After plating, the plate was centrifuged for 1min at 50rcf to collect organoids in the center of the wells. Organoids were visually assessed prior to the start of the assay and after its completion, only organoids entirely within the sensor at the center of the well in both assessments were included in the final analysis. To account for drug diffusion times, the time interval for initial readings and after FCCP and rotenone + antimycin A was increased to 36 minutes, while the interval after oligomycin was increased to 54 minutes with measurements taken every 6 mints. Additionally, the concentration of oligomycin was increased to 1.5uM. For normalization, individual organoids were collected after the assay’s completion and lysed in IX RIPA buffer (Sigma #R0278) for quantification with Precision Red (Cytoskeleton, Inc. #ADV02-A) according to the manufacturer’s protocol.

### CTPI-2 Treatment

For time lapse imaging of the CTIP-2 treatment assembloids were immobilized against a glass bottom dish as described above and their assembloid media was supplemented with 50uM CTPI-2 immediately prior to imaging. Assembloids were imaged and analyzed as described above. CTPI-2 toxicity was assessed by treating assembloids with 50uM CTPI-2 for 24hours prior to dissociation and staining with an Annexin V Cell Viability assay (Biolegend #422201 & #640935) according to the manufacturer’s protocol. Flow cytometry data was collected with a BD-Fortessa and analyzed with FlowJo v10.

