## Supplemental Figures for "Cortical assembloids support the development of fast-spiking human PVALB+ cortical interneurons and uncover schizophrenia-associated defects"

#### Supplemental Figure 1

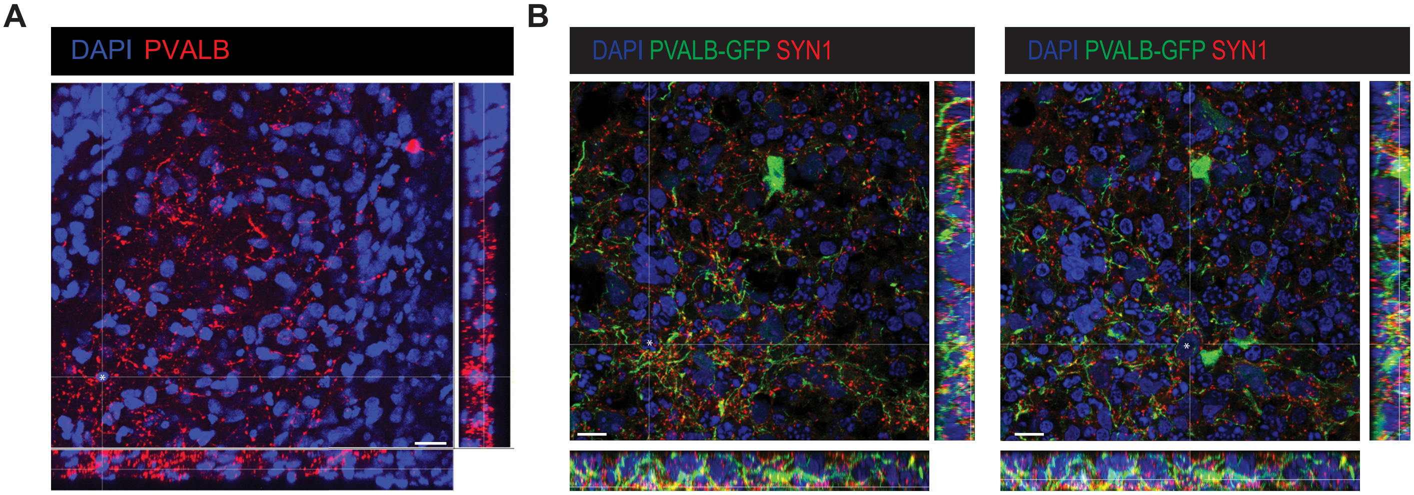

**Supplemental Figure 1 – Baskets and perisomatic boutons from PVALB+ cortical interneurons**. **A**) 3D stacks of PVALB staining on day 120 (left) and day 180 (right) showing perisomatic boutons surrounding PVALB- DAPI signals, indicating contact with the cell soma. Scale bar=20um.

**Supplemental Figure 2**

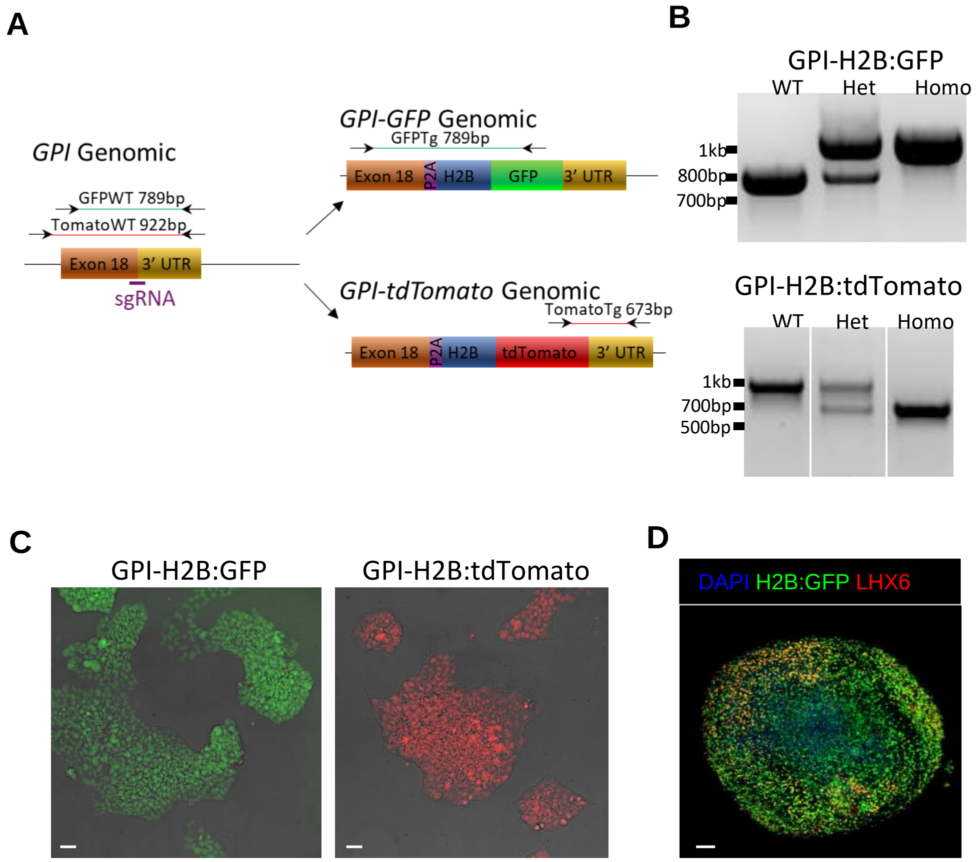

**Supplemental Figure 2– Generation of ubiquitous reporter lines for tracking and recovering cells from integrated neocortical organoids. A**) Schematic of the genome surrounding the last exon of the GPI locus (left). Location of sgRNA used along with genotyping primers and expected bands, color coded for either GFP or tdTomato genotyping, are shown above. The transgenic P2A-H2B:GFP and P2A-H2B:tdTomato are shown to the right along with genotyping primers and their respective expected bands. **B**) Genotyping PCR with the primers shown in panel A for GPI-H2B:GFP (Top) and GPI-H2B:tdTomato (Bottom). **C**) Undifferentiated ESCs targeted with either GPI-H2B:GFP (right) or GPI-H2B:tdTomato (Left). **D**) A d90 ventral forebrain organoid, note the maintained expression of nuclear GFP, particularly in LHX6+ cells.

### Supplemental Figure 3

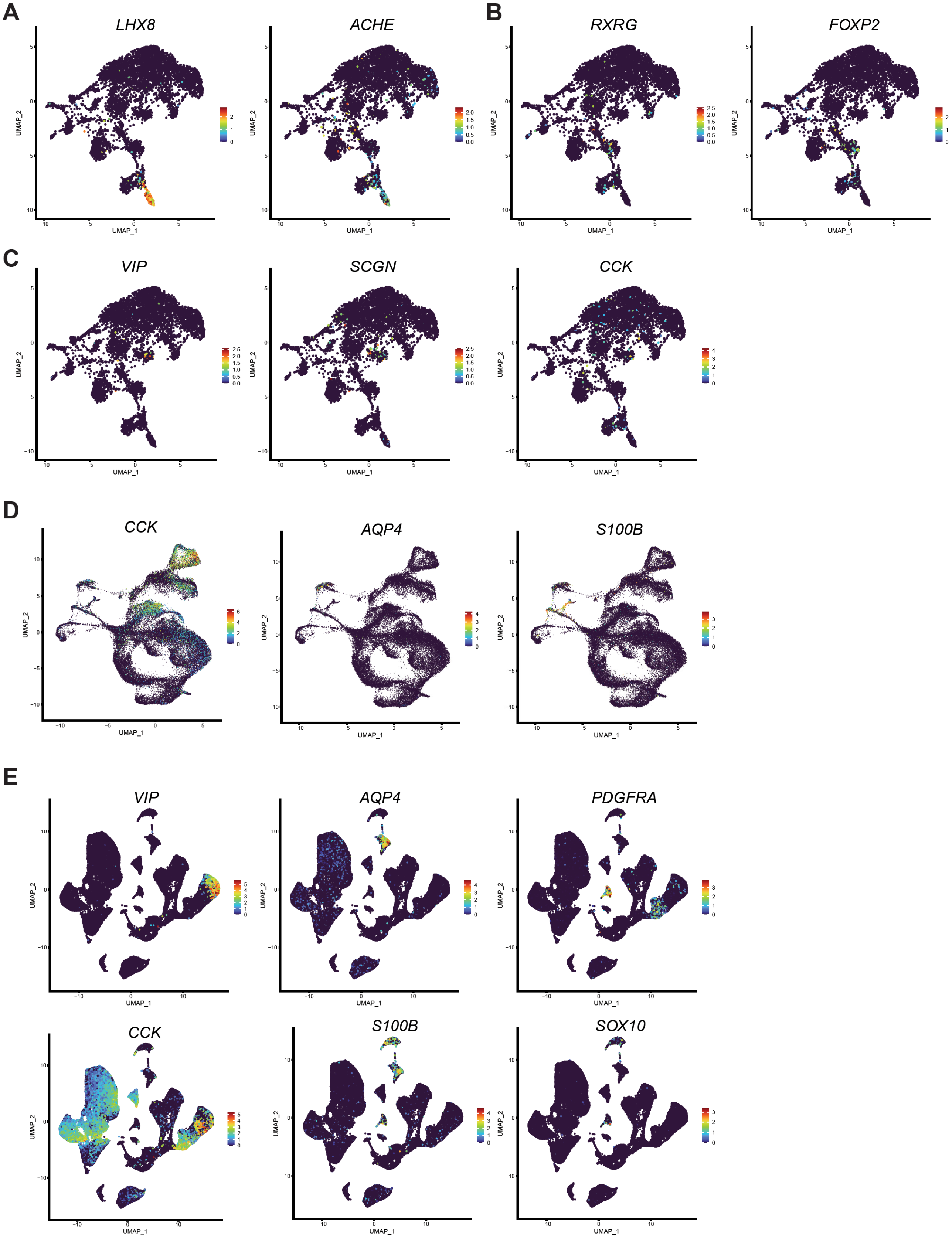

**Supplemental Figure 3. Subcortical and CGE derived neurons represent a minor population of vFB derived cells in cortical assembloids A**) Shown are select markers of MGE-derived striatal interneurons, they are predominantly coexpressed in cluster 14 (see **Fig. 3A** for clusters) **B**) Cluster 9 contains neurons positive for striatal and globus pallidus markers. **C**) CGE-derived cortical interneurons are evident in cluster 11. **D**) UMAP plots of integrated mouse and cortical assembloid samples related to figure 3B. Shown is an additional CGE-interneuron markers (CCK) and markers showing astrocytes in cluster 15 (AQP4 & S100B). **E**) UMAP plots of CGE markers cortical assembloid data integrated with postmortem human cortical data showing markers of CGE derived cortical interneurons (VIP & CCK), astrocytes (AQP4 & S100B), and oligodendrocyte precursor cells (SOX10 & PDGFRB).

**Supplemental Figure 4**

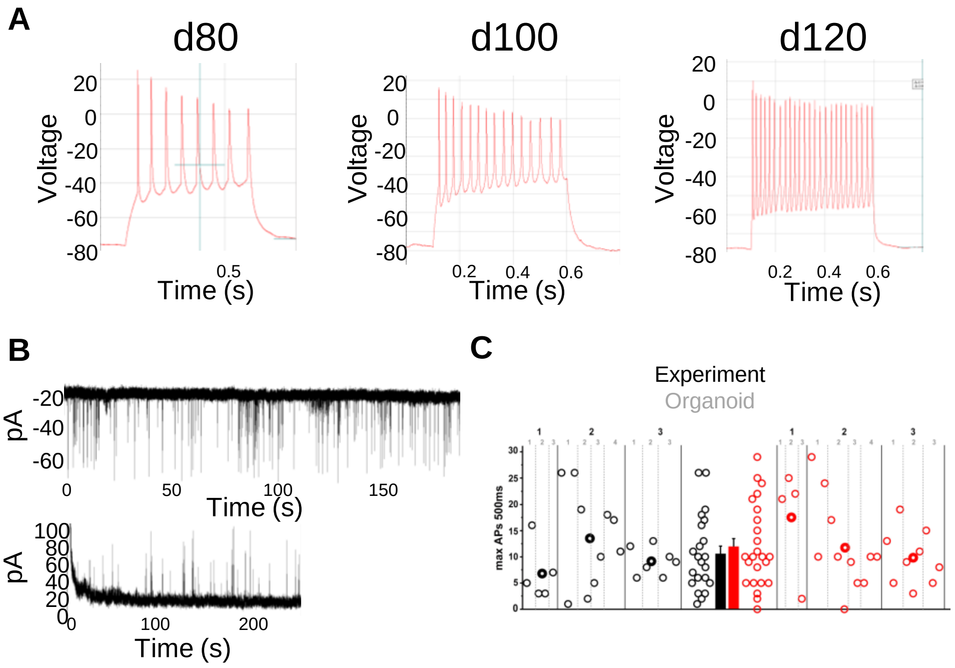

**Supplemental Figure 4 – Comparison electrophysiologic properties of 22q11.2 deletion and isogeneic control cortical interneurons at day 120. A**) Representative traces of action potentials recorded from CINs at days 80, 100 and 120, showing evidence of maturation proceeding over time. **B**) Representative traces of CINs displaying spontaneous synaptic activity at day 100. Neurons were held at -70mV to detect glutamatergic (top) and 0mV for GABAergic (bottom) synaptic events, showing robust synaptic activity. **C**) Quantification of maximum action potentials recorded per cell from 3 independent experiments at day 120. The experiment is indicated along the top Y-axis in black, while the individual organoid from that experiment is indicated right below that in gray, each circle represents an individual CIN. The mean with s.e.m. is plotted to the right side of the graph. **C**) Quantification of max APs for WT neurons (left, black) and 22q11.2 deletion neurons (right, red), recorded at day 120 with GPI-GFP and GPI-tdTomato reporters.

### Supplemental Figure 5

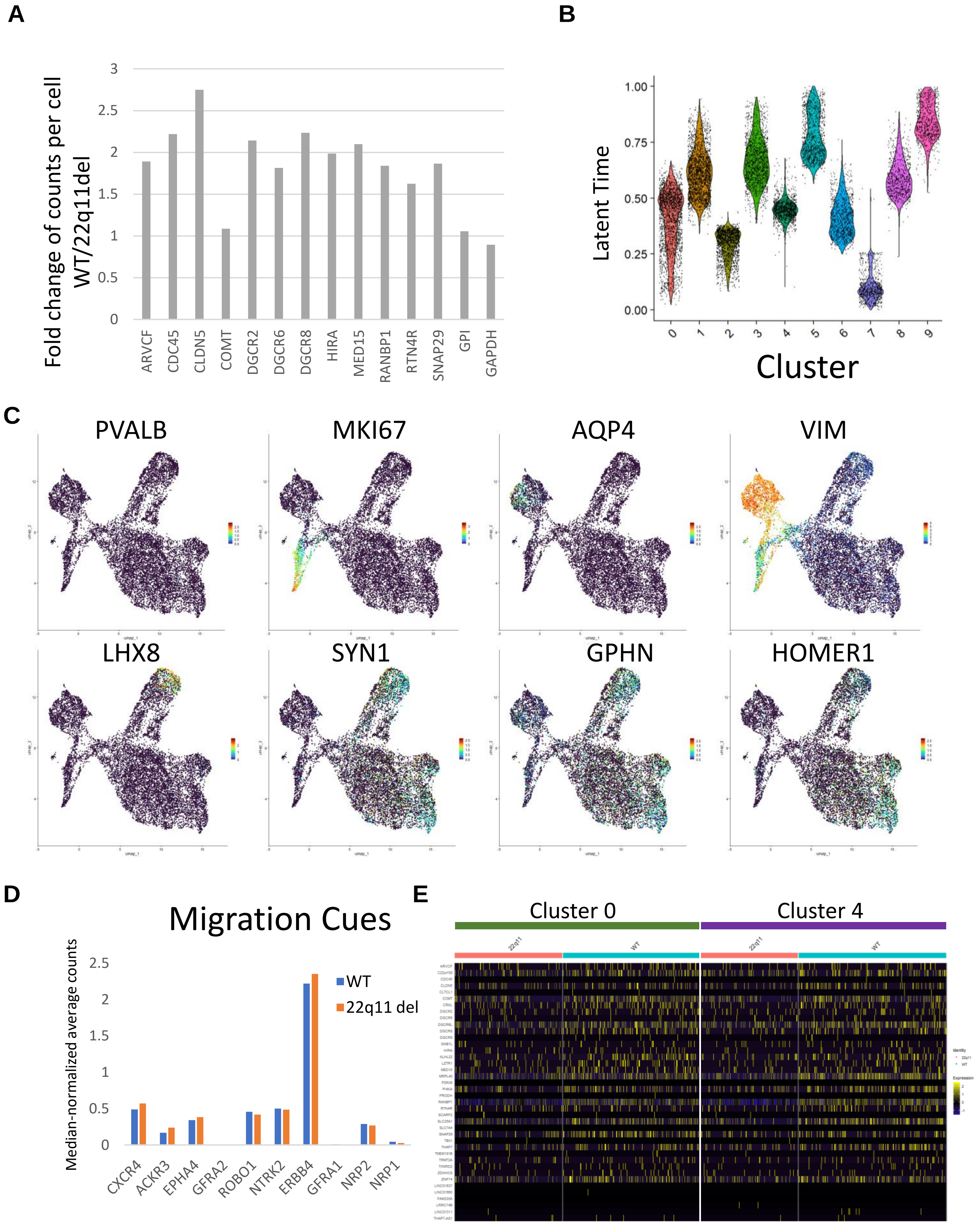

**Supplemental Figure 5**. **Molecular comparison of 22q11.2 deletion cortical interneurons to isogenic controls.** **A**) Global fold change values (WT/22q11.2del) for select genes within the 22q11.2 locus. Housekeeping genes GPI and GAPDH are not in the 22q11.2 locus and included as a control. **B**) Latent times calculated from scVelo are shown for each cluster. **C**) Additional gene expression data from the 22q11.2 deletion experiment relevant to Fig. 6A. **D**) No difference in expression of genes known to be involved in cortical interneuron migration guidance. Shown are normalized count values between 22q11.2 deletion and control neurons in the intermediate migrating population (clusters 0 & 4). **E**) Heatmap showing expression of all transcripts within the 22q11.2 locus detected in the scRNAseq experiments, grouped by cluster and genotype.

**Supplemental Figure 6**

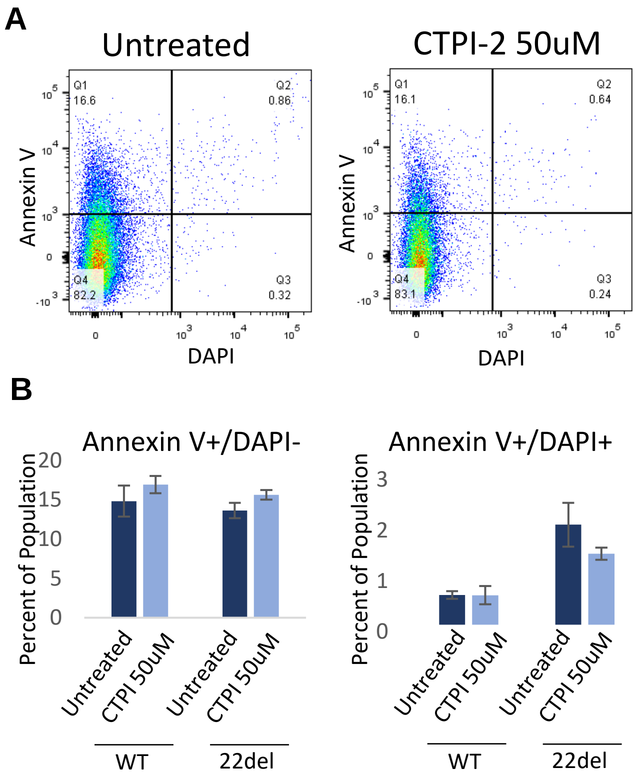

**Supplemental Figure 6. CTIP-2 treatment does not induce cell death in vFB organoids. A**) Representative flow plots of annexin V stain of vFB organoids after 24h treatment with 50uM CTPI-2. **B**) Quantification of A. n=3 organoids per treatment group. * = $p<0.05$; ****= $p<0.0001$. Error bars = s.e.m.

#### Supplemental Figure 7

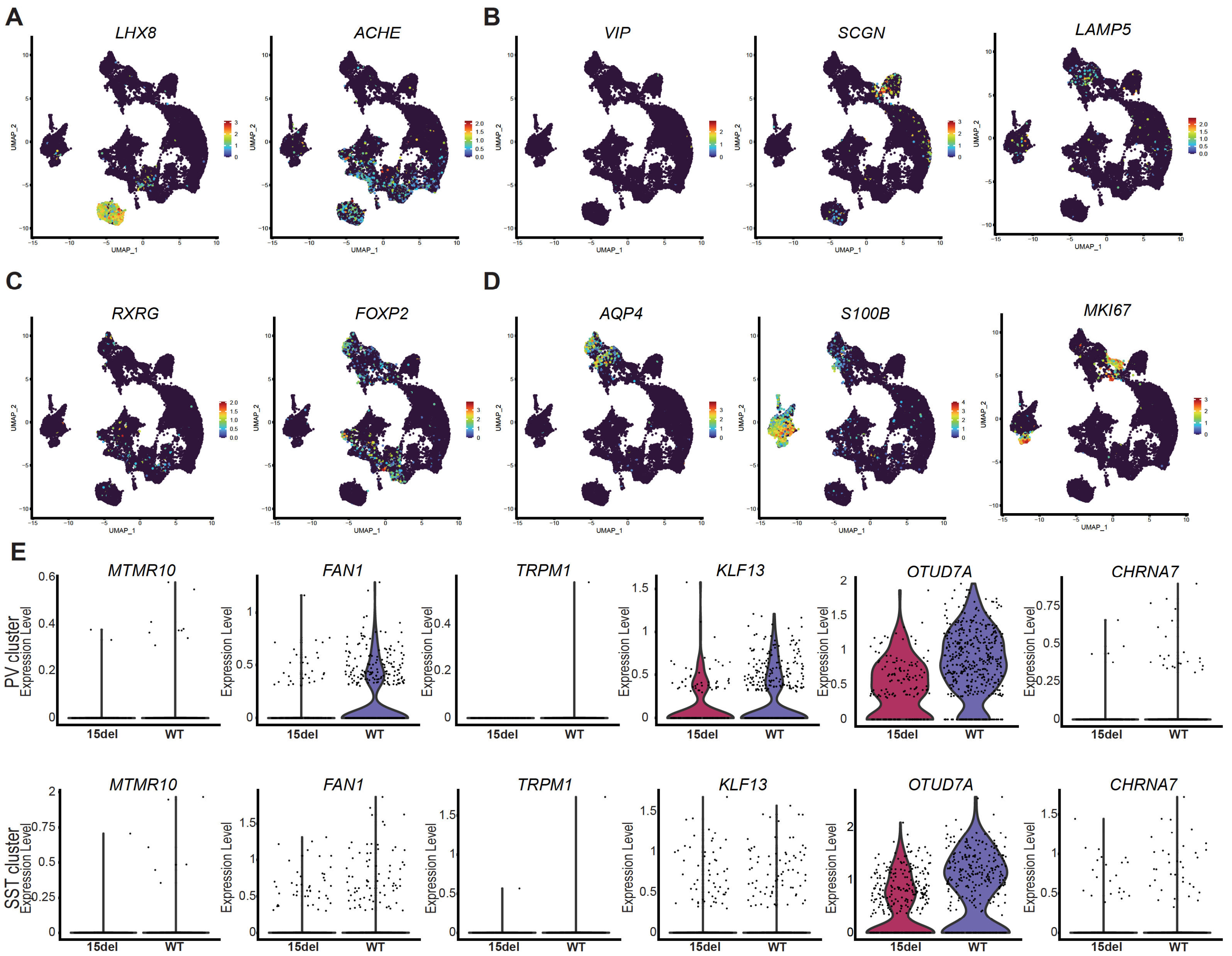

**Supplemental Figure 7. Additional molecular analysis of day 125 15q13 deletion organoids. A)** Markers demonstrating populations of MGE derived subcortical interneurons. **B)** CGE-derived interneuron markers. **C**) Striatal and globus pallidus neuron markers. **D**) Markers of astrocytes (AQP4 & S100B) and cycling progenitors (MKI67). **E**) Differential expression of genes within the 15q13 locus in the PVALB cluster (top) and SST cluster (bottom).

**Legends to Supplementary Videos:**

**Supplemental Video 1. vFB derived cells migrate throughout the dFB domain of the cortical assembloid**. A representative example of a day 60 assembloid with the GPI-H2B:tdTomato labeled vFB shown in red and the GPI-H2B:GFP labeled dFB shown in green.

**Supplemental Video 2. Migration of cortical interneurons in cortical assembloids.** Representative time lapse imaging of GPI-cytoplasmic GFP labeled cortical interneurons migrating into the cortical assembloid at day 35 after being aggregated at day 30. An approximately 62um Z-section was acquired every 20 minutes for 14 hours.

**Supplemental Video 3. Migration of SV lines and isogenic controls throughout the cortical assembloids.** Shown are representative examples of iDISCO stained and cleared cortical assembloids at day 60 displaying cortical interneuron migration throughout the cortical assembloid. The dFB portion of the assembloid was unlabeled, while the vFB was generated with SV and isogenic control lines alternately labeled with either GFP or tdTomato.

**Supplemental Video 4. Migration of WT and 22q11.2 deletion cortical interneurons through the cortical assembloid.** H2B:tdTomato 22q11.2 cortical interneurons and H2B:GFP isogenic control cortical interneurons migrate through the dFB region of the cortical assembloid.
