## Supplementary material for "Cortical assembloids support the development of fast-spiking human PVALB+ cortical interneurons and uncover schizophrenia-associated defects": Table 1

| Structural Variant | Odds Ratio | p-value | Genes Affected | Size | Deletion Nature | CIN Phenotype in animal models | Reference |
|---|---|---|---|---|---|---|---|
| 22q11.2 Deletion | 15.6-∞ | $7.3\times10^{-13}$ | COMT, DGCR8 & 42 others | 2.52mb | NAHR | Impaired CIN migration into the cortex | Murphy et al. 1999; Kirov et al. 2009; Levinson et al. 2011; Meechan et al. 2012 |
| 15q13.3 deltion | 11.54-17.9 | 0.0029-$5.3\times10^{-4}$ | CHRNA7, OTUD7A & 5 others | 1.58mb | NAHR | Reduced CIN firing rate in mouse model | Stefansson et al. 2008; International Schizophrenia Consortium 2008; Nilsson et al. 2016 |
| 2q34 deletion* | 5.388* | 0.30* | ERBB4 | 400kb | NHEJ | Severely reduced CINs in cortex, failure in migration of CINs to cortex, reduced synapses and excitation of PVALB+ CINs | Walsh et al. 2008; Flames et al. 2004; Krivosheya et al. 2008; Fazzari et al. 2010 |

**Genetic lesions associated with schizophrenia to be studied in this proposal**.
*2q34 was a *de novo* mutation detected in a patient with childhood onset schizophrenia and not statistically significant. NAHR: Nonallelic Homologous Recombination; NHEJ: Nonhomologous End Joining
